# Liraglutide exerts protective effects on MASLD independently of lipophagy, and the combination with kombucha presents no additional advantage

**DOI:** 10.1101/2025.10.24.684466

**Authors:** Jenifer Souza de Almeida, Sandra Andreotti, Tatiana Yoshimi Obara, Bianca Lugatti Cardoso Ferreira, Rodrigo dos Santos Ono Gentile, Felipe Nunes de Camargo, Ayumi Cristina Medeiros Komino, Carla Roberta de Oliveira Carvalho

**Author notes:** Correspondence (C.R.O.C.). (J.S.A.); (S.A.); (T.Y.O.); (B.L.C.F.); (R.S.O.G.); (F.N.C.); (A.C.M.K.).

## Abstract

Metabolic dysfunction – associated steatotic liver disease (MASLD) has a high global prevalence, requiring effective therapeutic options. This study investigated the effects of liraglutide and kombucha, alone and in combination, in a murine model of high-fat diet (HFD)-induced MASLD. Metabolic parameters, body composition, liver histology, lipid profile, and markers of lipophagy were evaluated. The results demonstrate that liraglutide reduced body weight, fat mass, and lean mass, as well as food and caloric intake, fasting blood glucose, hepatic triglycerides (TG), steatosis, and fibrosis, improved the lipid profile and glucose tolerance, and increased lysosomal acid lipase (LAL) expression in the liver. Kombucha reduced lean mass, liver steatosis, hepatic TG, food and caloric intake, fasting blood glucose, low-density lipoprotein cholesterol (LDL-C), and liver fibrosis, in addition to improving glucose tolerance. However, the combination of treatments did not show synergistic effects superior to those of liraglutide alone. Although liraglutide independently impacted LAL expression, none of the other lipophagy markers were modulated in the liver, suggesting that the beneficial mechanisms of action are independent of the lipophagy pathway. It is concluded that both liraglutide and kombucha are valid strategies for mitigating MASLD, but the combination does not offer superior advantages over liraglutide monotherapy.

## 1. Introduction

Metabolic dysfunction-associated steatotic liver disease (MASLD) is characterized by a spectrum of progressive liver changes, with hepatic steatosis (ectopic fat accumulation ≥5% in hepatocytes) being its initial stage, in which there is no significant hepatocellular inflammation. On the other hand, the later stage of MASLD, known as metabolic dysfunction-associated steatohepatitis (MASH), is characterized by the presence of hepatocellular inflammation and an increased risk of developing fibrosis, cirrhosis, and/or hepatocellular carcinoma, with a consequent higher mortality risk [1,2].

The continued rise in the adoption of a sedentary lifestyle, currently observed, favors the increase in the global prevalence of obesity and, concomitantly, the prevalence of MASLD, which reaches a quarter of the world’s adult population [3]. The increasing global impact of MASLD warrants attention, particularly due to its socioeconomic implications and health consequences. In the United States, this disease is among the main causes of hepatocellular carcinoma, resulting in the need for liver transplantation. In this context, it is worth noting that in 2016, the lifetime costs of medical care provided to patients with MASLD in this country alone were approximately US$100 billion [4].

Some studies suggest that obesity and MASLD may impair the occurrence of lipophagy, an autophagic process that selectively targets lipid vesicles, thereby contributing to increased hepatic lipid accumulation and progression of MASLD [5,6]. In lipophagy, lipid droplets interact with cellular structures called autophagosomes, which subsequently fuse with lysosomes, enabling the degradation of lipid content by lysosomal acid lipase (LAL) [7,8]. This process can be summarized into five steps (1) formation of a double-membrane structure called phagosome from the endoplasmic reticulum, (2) nucleation and expansion of the phagosome, (3) maturation of this into an autophagosome and interaction with lipid droplets, (4) fusion of autophagosome with lysosome, (5) and release of lipid droplets into the lysosomal lumen for hydrolysis mediated by LAL [7,9].

In the therapeutic context, the impact of pharmacological and non-pharmacological agents on MASLD is under investigation. Previous results in the literature and those observed in our laboratory indicate that liraglutide, a glucagon-like peptide-1 (GLP-1) receptor agonist and a drug approved by the Food and Drug Administration (FDA) for the treatment of type 2 diabetes mellitus (T2DM), has a beneficial effect on MASLD in murine models. Indeed, besides pharmacological agents, probiotics have been shown to have therapeutic potential for MASLD and could potentially be a more accessible and lower-cost therapeutic option for the population [10]. Interestingly, kombucha, a probiotic beverage, has been shown to offer a hepatoprotective effect against liver damage in experimental models of hepatic steatosis [11-13]. Additionally, our group demonstrated that supplementation with this drink for only 9 days was sufficient to promote improvements in hepatic steatosis, glucose tolerance, hyperinsulinemia, and hepatic collagen fiber deposition in a model of MASLD induced by a high-fat diet (HFD) [14].

In the present study, we hypothesized that the concomitant use of liraglutide and kombucha could amplify the effect of either alone on HFD-induced MASLD, and that this effect was related to lipophagy in the hepatocytes of this experimental model.

## 2. Results

### 2.1 Liraglutide reduces body weight and fat content in obese animals

As expected, HFD consumption was effective in increasing body weight after 10 weeks of diet (Figure S1, A - C: 29,825 ± 2,648 vs. L–/K–: 39,298 ± 4,761 g; P = 0.0026), increasing the area under the curve (AUC) of body weight over the final 2 weeks of the protocol (Figure S1, B and C - C: 418,500 ± 35,406 vs. L–/K–: 567,600 ± 66,317 g.day; P = 0.0010), final body weight (Figure S1, D - C: 28,940 ± 2,119 vs. L–/K–: 40,127 ± 5,140 g; P = 0.0004), and lean mass content (Figure S1, E - C: 19,205 ± 1,234 vs. L-/K-: 22,695 ± 2,556 g; P = 0.0229) and fat mass content (Figure S1, F - C: 2,603 ± 0,359 vs. L-/K-: 10,869 ± 1,380g; P < 0.0001) in animals. Animals treated with or without liraglutide and kombucha started the treatment protocol with equal body weight [Figure 1, A - L factor: F (1, 40) = 0.001767, P = 0.9667; K factor: F (1, 40) = 0.1789, P = 0.6746; Interaction: F (1, 40) = 0.01607, P = 0.8998], and throughout the treatment period, the effect of liraglutide-containing regimens was distinct from that of regimens without the drug in terms of AUC of body weight [Figure 1, B and C - L factor: F (1, 40) = 13.60, P = 0.0007; K factor: F (1, 40) = 1.594, P = 0.2141; Interaction: F (1, 40) = 1.187, P=0.2825] and final body weight [Figure 1, D - L factor: F (1, 40) = 26.71, P < 0.0001; K factor: F (1, 40) = 2.640, P = 0.1121; Interaction: F (1, 40) = 2.861, P = 0.0985], which was not observed with kombucha. Similarly, no significant interaction between factors was observed in these parameters, further reinforcing the impact of liraglutide on weight dynamics.

**Figure 1.**
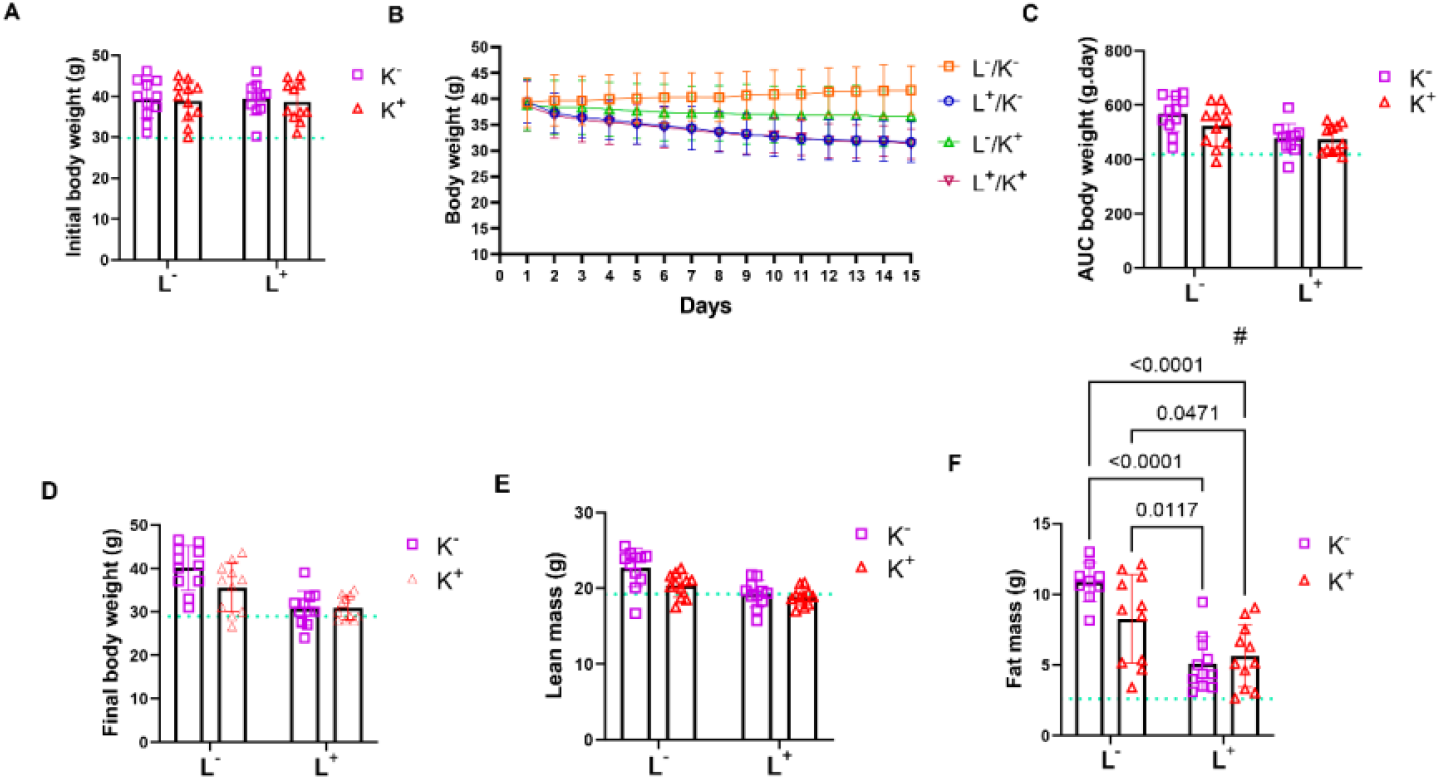
Body weight and composition. The results were obtained through a Two-Way ANOVA and are presented as mean ± standard deviation (SD), with a significance level of P ≤ 0.05. L factor = liraglutide factor; K factor = kombucha factor; interaction = interaction between factors. The results of the multiple comparison test are presented as P values, indicating significant interactions between factors and significant differences between groups, as noted in each item. The green line indicates the MD±SD of group C, and each scatter plot indicates a sampling unit. # indicates a significant interaction between liraglutide and kombucha. (**A**) Initial body weight, obtained before the start of treatments (n= 11 in all groups). Two-Way ANOVA: L factor - F (1, 40) = 0.001767, P=0.9667; K factor - F (1, 40) = 0.1789, P=0.6746; Interaction - F (1, 40) = 0.01607, P=0.8998. (**B**) Time course of body weight from the 1st to the 15th day of treatment (n=11 in all groups). (C) Area under the curve (AUC) of body weight of animals throughout the treatment period (n=11 in all groups). Two-Way ANOVA: L factor - F (1, 40) = 13.60, P=0.0007; K factor - F (1, 40) = 1.594, P=0.2141; Interaction - F (1, 40) = 1.187, P=0.2825. (**D**) Final body weight, obtained after the last day of the treatment period (n= 11 in all groups). Two-Way ANOVA: L factor - F (1, 40) = 26.71, P<0.0001; K factor - F (1, 40) = 2.640, P=0.1121; Interaction - F (1, 40) = 2.861, P=0.0985. (E) Lean body mass content, obtained after the last day of the treatment period (n=11 in all groups). Two-Way ANOVA: L factor - F (1, 40) = 19.93, P<0.0001; K factor - F (1, 40) = 6.062, P=0.0182; Interaction - F (1, 40) = 3.068,P=0.0875. (F) Fat body mass content, obtained after the last day of the treatment period (n=9 in L-/K-, and n=11 in the other groups). Two-Way ANOVA: L factor - F (1, 38) = 35.46, P<0.0001; K factor - F (1, 38) = 2.109, P=0.1546; Interaction - F (1, 38) = 4.932, P=0.0324.

Both liraglutide [Figure 1, E - L factor: F (1, 40) = 19.93, P < 0.0001] and kombucha [Figure 1, E -K factor: F (1, 40) = 6.062, P = 0.0182] factors showed a significant effect on lean mass content, but there was no significant interaction between such factors [Figure 1, E - Interaction: F (1, 40) = 3.068, P = 0.0875], indicating that they possibly act independently on this parameter.

Although the presence of liraglutide modulated fat mass content [Figure 1, F - L factor: F (1, 38) = 35.46, P < 0.0001] and that of kombucha did not [Figure 1, F - K factor: F (1, 38) = 2.109, P=0.1546], there was a significant interaction between these factors [Figure 1, F - Interaction: F (1, 38) = 4.932, P=0.0324]. In this sense, the combination of treatments was effective in reducing fat mass in animals fed with HFD (Figure 1, F - L+/K+: 5,637 ± 2,201 vs. L-/K-: 10,869±1,380 g; P < 0.0001), an effect that was not distinct from that promoted by liraglutide (Figure 1, F - L+/K+: 5,637 ± 2,201 vs. L+/K-: 5,095±1,901 g; P = 0.9437 and Figure 1, F - L+/K-: 5,095±1,901 vs. L-/K-: 10,869 ± 1,380 g; P < 0.0001). Kombucha did not alter the fat mass content of obese animals (Figure 1, F - L-/K+: 8,275 ± 3,126 vs. L-/K-: 10,869 ± 1,380 g; P = 0.0710). However, the combination of treatments resulted in a lower fat mass content compared to animals treated with kombucha alone (Figure 1, F - L+/K+: 5,637± 2,201 vs. L-/K+: 8,275 ± 3,126 g; P = 0.0471), as did liraglutide (Figure 1, F - L+/K-: 5,095±1,901 vs. L- /K+: 8,275 ± 3,126 g; P = 0.0117).

### 2.2. Liraglutide and kombucha reduce food and caloric intake in animals fed HFD

Animals fed the HFD showed higher caloric intake compared to control animals throughout the 12 consecutive weeks of feeding (Table S1 - C: 677,560 ± 22,335 vs. L-/K-: 854,933 ± 101,328 kJ.day; P = 0.0069), although food intake was lower in this group (Table S1 - C: 50,608 ± 1,671 vs. L-/K-: 38,195 ± 4,524 g.day; P = 0.0003). Both factors, liraglutide [Table 1 - L factor: F (1, 23) = 73.44, P < 0.0001) and kombucha [Table 1 - K factor: F (1, 23) = 10.95, P = 0.0031], affected the food intake of obese animals, and the interaction between these factors was significant [Table 1 - Interaction: F (1, 23) = 12.30, P = 0.0019]. In this regard, the combination of treatments reduced the AUC of food intake in animals fed HFD (Table 1 - L+/K+: 20,589 ± 4,524 vs. L-/K-: 38,195 ± 4,524 g.day; P < 0.0001), having greater efficacy than kombucha (Table 1 - L+/K+: 20,589± 4,524 vs. L-/K+: 28,093 ± 3,076 g.day; P = 0.0067 and Table 1 - L-/K+: 28,093 ± 3,076 vs. L-/K-: 38,195 ± 4,524 g.day; P = 0.0005), but not liraglutide (Table 1 - L+/K+: 20,589 ± 4,524 vs. L+/K-: 20,296 ± 3,099 g.day; P = 0.9989 and Table 1 - L+/K-: 20,296 ± 3,099 vs. L-/K-: 38,195 ± 4,524 g.day; P < 0.0001).

**Table 1.** Food and caloric intake.

| Group | AUC Food intake (g.day)<br>Mean ± SD | AUC Caloric intake (kJ.day)<br>Mean ± SD |
| --- | --- | --- |
| L-/K- | 38,195 ± 4,524 | 854,933 ± 101,328 |
| L+/K- | 20,296 ± 3,099**** | 454,257 ± 69,328**** |
| L-/K+ | 28,093 ± 3,076***\$\$ | 628,814 ± 68,842***\$\$ |
| L+/K+ | 20,589 ± 4,524****&& | 460,857 ± 101,297****&& |

Simultaneously, liraglutide [Table 1 - L factor: F (1, 23) = 73.42, P < 0.0001] and kombucha [Table 1 - K factor: F (1, 23) = 10.94, P = 0.0031] had a significant effect on caloric intake, with a significant interaction between these factors [Table 1 - Interaction: F (1, 23) = 12.30, P = 0.0019]. The combination of liraglutide and kombucha reduced the AUC of caloric intake in obese animals (Table 1 - L+/K+: 460,857 ± 101,297 vs. L-/K-: 854,933 ± 101,328 kJ.day; P < 0.0001), and this effect was superior to that of kombucha (Table 1 - L+/K+:460,857 ± 101,297 vs. L-/K+:628,814 ± 68,842 kJ.day; P = 0.0068 and Table 1 - L-/K+:628,814 ± 68,842 vs. L-/K-:854,933 ± 101,328 kJ.day; P = 0.0005), but not to that of liraglutide (Table 1 - L+/K+: 460,857 ± 101,297 vs. L+/K-: 454,257 ± 69,328 kJ.day; P = 0.9989 and Table 1 - L+/K-: 454,257 ± 69,328 vs. L-/K-: 854,933 ± 101,328 kJ.day; P < 0.0001). These data reinforce the effect of the drug liraglutide on regulating food intake and, consequently, caloric intake.

The results were obtained through Two-way ANOVA and are presented as MD ± SD. The significance level adopted was P ≤ 0.05. L factor = liraglutide factor; K factor = kombucha factor; interaction = interaction between factors. **** indicates P ≤ 0.0001 in relation to L-/K-, *** indicates P ≤ 0.001 in relation to L-/K-, $$ indicates P ≤ 0.01 in relation to L+/K-, and && indicates P ≤ 0.01 in relation to L-/K+. AUC of food intake during the treatment period (n = 6 in the L-/K- group, and n = 7 in the other groups). Two-Way ANOVA: L factor - F (1, 23) = 73.44, P < 0.0001; K factor - F (1, 23) = 10.95, P = 0.0031; Interaction - F (1, 23) = 12.30, P = 0.0019. AUC of caloric intake throughout the treatment period (n = 6 in the L-/K- group, and n = 7 in the other groups). Two-Way ANOVA: L factor - F (1, 23) = 73.42, P < 0.0001; K factor - F (1, 23) = 10.94, P = 0.0031; Interaction - F (1, 23) = 12.30, P = 0.0019.

### 2.3 Liraglutide and kombucha reduce fasting blood glucose and improve oral glucose tolerance in animals with diet-induced obesity

In addition to the effects on the weight and feeding dynamics of the animals, HFD consumption resulted in increased fasting blood glucose (Figure S2, A - C: 7,236 ± 0,606 vs. L-/K-: 9,096 ± 1,479 mmol.L^-1^; P = 0.0323) and reduced oral glucose tolerance (Figure S2, B and C - C: 1212,800 ± 184,901 vs. L-/K-: 1653,000 ± 330,207 mmol.L^-1^.min; P = 0.0153). The liraglutide factor showed an impact on fasting blood glucose [Figure 2, A - L factor: F (1, 39) = 33.77, P < 0.0001], in contrast to the kombucha factor [Figure 2, A - K factor: F (1, 39) = 3.600, P = 0.0652]. The interaction between both factors was significant [Figure 2, A - Interaction: F (1, 39) = 4.299, P = 0.0448]. The combination of treatments was effective in reducing fasting blood glucose in the animals (Figure 2, A - L+/K+: 6,596 ± 0,570 vs. L-/K-: 9,096±1,479 mmol.L^-1^; P < 0.0001), however, its efficacy was not superior to that of liraglutide (Figure 2, A - L+/K+: 6,596±0,570 vs. L+/K-: 6,539 ± 0,634 mmol.L^-1^; P = 0.9993 and Figure 2, A - L+/K-: 6,539 ± 0,634 vs. L-/K-: 9,096 ± 1,479 mmol.L^-1^; P < 0.0001) or that of kombucha (Figure 2, A - L+/K+: 6,596 ± 0,570 vs. L-/K+: 7.808 ± 1.236 mmol.L^-1^; P = 0.0508 and Figure 2, A - L-/K+: 7.808 ± 1.236 vs. L-/K-: 9.096±1.479 mmol.L^-1^; P = 0.0343). These data indicate that the effect of kombucha on fasting blood glucose is dependent on the action of liraglutide, highlighting the impact of this drug on blood glucose regulation.

**Figure 2.**
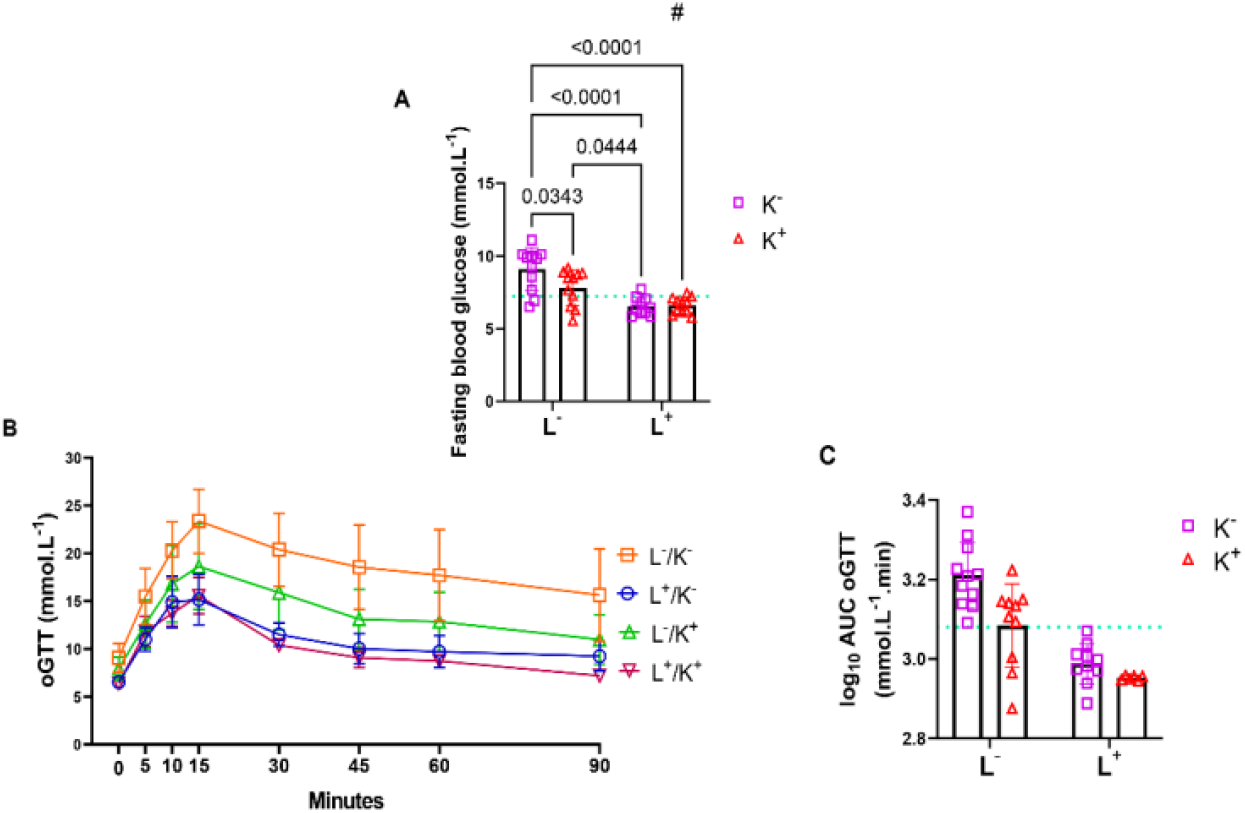
Fasting blood glucose and glucose tolerance. The results were obtained through Two-wayANOVA and are presented as MD±SD. The significance level adopted was P ≤ 0.05. L factor = liraglutide factor; K factor = kombucha factor; interaction = interaction between factors. The results of the multiple comparison test are presented as P values, indicating significant interactions between factors and significant differences between groups, as noted in each item. The green line indicates the MD±SD of group C, and each scatter plot indicates a sampling unit. # indicates a significant interaction between liraglutide and kombucha. (**A**) Fasting blood glucose levels obtained at the end of the treatment period (n = 10 in the L+/K- group, and n = 11 in the other groups). Two-Way ANOVA: L factor - F (1, 39) = 33.77, P < 0.0001; K factor - F (1, 39) = 3.600, P = 0.0652; Interaction - F (1, 39) = 4.299, P = 0.0448. (**B**) Time course of the oral glucose tolerance test (oGTT) performed at the end of the treatment period (n = 11 in the L-/K-, L-/K+, L+/K+ groups, and n = 10 in L+/K-). (C) AUC of oGTT transformed into base-10 logarithm (log_10_) (n= 11 in the L-/K- group, n = 10 in L-/K+ and L+/K-, and n = 6 in L+/K+). Two-Way ANOVA: L factor - F (1, 33) = 47.67, P < 0.0001; K factor - F (1, 33) = 9.998, P = 0.0034. Interaction - F (1, 33) = 3.199, P = 0.0829.

Although the factors liraglutide [Figure 2, B and C - L factor: F (1, 33) = 47.67, P < 0.0001] and kombucha [Figure 2, B and C - K factor: F (1, 33) = 9.998, P = 0.0034] showed an effect on the AUC of the oral glucose tolerance test (oGTT), there was no significant interaction between them, indicating that the treatments may act through independent pathways [Figure 2, B and C - Interaction: F (1, 33) = 3.199, P = 0.0829].

### 2.4. Liraglutide improves plasma lipids, and kombucha reduces plasma low-density lipoprotein cholesterol (LDL-C) levels in animals with HFD-induced obesity

Animals fed HFD showed significant increases in plasma levels of triglycerides (TG) (Figure S3, A- C: 0.484 ± 0.083 vs. L-/K-: 0.755 ± 0.193 mmol.L^-1^; P = 0.0191), total cholesterol (TC) (Figure S3, B - C: 2.212 ± 0.186 vs. L-/K-: 3.652 ± 0.584 mmol.L^-1^; P = 0.0006), high-density lipoprotein cholesterol (HDL-C) (Figure S3, C - C: 1.784 ± 0.219 vs. L-/K-: 2.591 ± 0.506 mmol.L^-1^; P = 0.0106), LDL-C (Figure S3, D - C: 0.331 ± 0.094 vs. L-/K-: 1.222 ± 0.481mmol.L^-1^; P = 0.0054) and very low density lipoprotein cholesterol (VLDL-C) (Figure S3, E - C: 0.097 ± 0.016 vs. L-/K-: 0.151 ± 0.039mmol.L^-1^; P = 0.0198) at the end of the experimental protocol. The presence of liraglutide significantly modulated plasma TG levels [Figure 3, A - L factor: F (1, 38) = 19.40, P < 0.0001], which was not observed with the presence of kombucha [Figure 3, A - K factor - F (1, 38) = 2.487, P = 0.1231]. The interaction between the factors was not significant [Figure 3, A - Interaction: F (1, 38) = 0.3888, P = 0.5367]. The same was observed for plasma levels of TC [Figure 3, B - L factor: F (1, 38) = 18.92, P < 0.0001; K factor: F (1, 38) = 0.0006980, P=0.9791; Interaction: F (1, 38) = 0.1338, P = 0.7166), HDL-C [Figure 3, C - L factor: F (1, 39) = 5.330, P=0.0263; K factor: F (1, 39) = 0.2944, P = 0.5905; Interaction: F (1, 39) = 0.8297, P = 0.3680] and VLDL-C [Figure 3, E - L factor: F (1, 38) = 19.21, P < 0.0001; K factor: F (1, 38) = 2.471, P = 0.1243; Interaction: F (1, 38) = 0.4009, P = 0.5304]. In contrast, liraglutide and kombucha affected LDL-C, with the interaction between them being significant [Figure 3, D - L factor: F (1, 30) = 115.6, P < 0.0001; K factor: F (1, 30) = 6.529, P = 0.0159; Interaction: F (1, 30) = 7.095, P = 0.0123]. Therefore, the combination of treatments was effective in reducing plasma LDL-C in obese animals (Figure 3, D - L+/K+: -0.378 ± 0.053 vs. L-/K-: 0.062 ± 0.157 mmol.L^-1^; P < 0.0001), which was superior to that of kombucha (Figure 3, D - L+/K+: ≤0.378 ± 0.053 vs L-/K+: ≤0.110±0.109 mmol.L^-1^; P < 0.0001 and Figure 3, D - L-/K+: “0.110 ± 0.109 vs. L- /K-: 0.062 ± 0.157 mmol.L^-1^; P = 0.0101), but not to that of liraglutide (Figure 3, D - L+/K+: “0.378 ± 0.053 vs. L+/K-: “0.381 ± 0.062 mmol.L^-1^; P = 0.9998 and Figure 3, D - L+/K-: “0.381 ± 0.062 vs. L- /K-: 0.062 ± 0.157 mmol.L^-1^; P < 0.0001). Liraglutide also showed greater efficacy than kombucha (Figure 3, D - L+/K-: “0.381 ± 0.062 vs. L-/K+: “0.110 ± 0.109 mmol.L^-1^; P < 0.0001).

**Figure 3.**
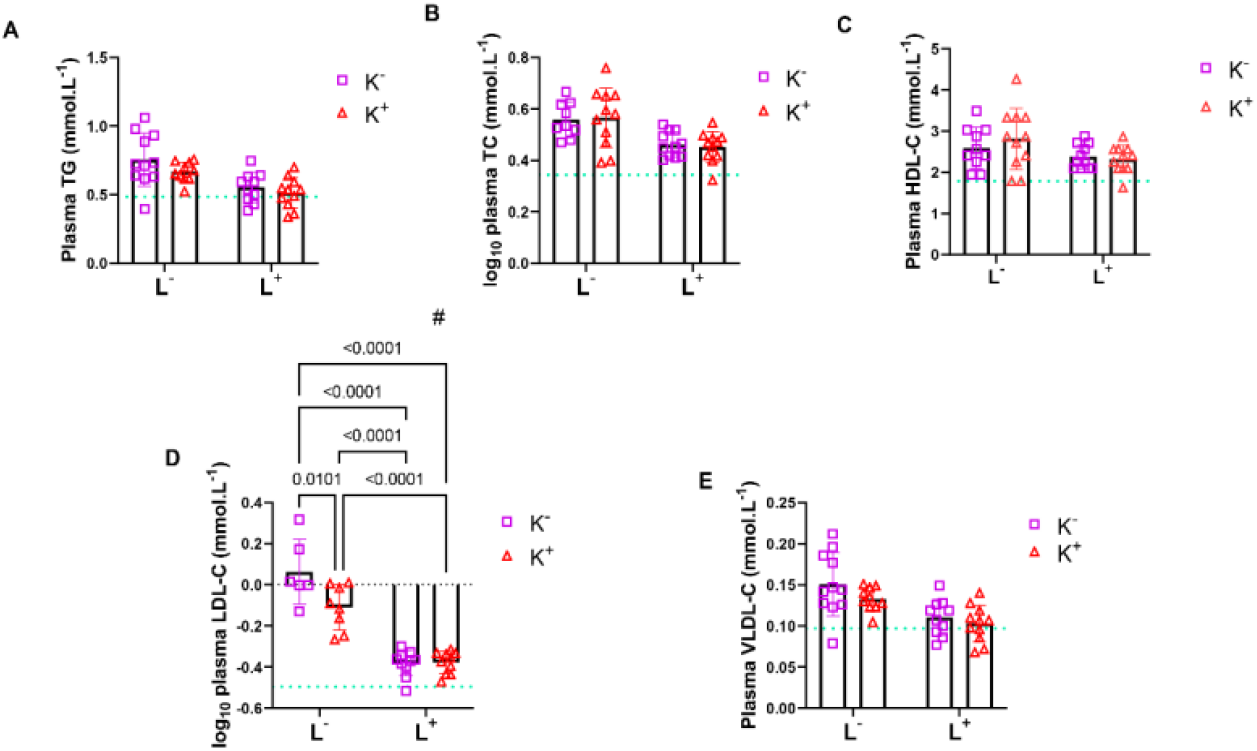
Plasma lipids. The results were obtained through a Two-Way ANOVA and are presented as mean ± standard deviation (SD), with a significance level of P ≤ 0.05. L factor = liraglutide factor; K factor = kombucha factor; interaction = interaction between factors. The results of the multiple comparison test are presented as P values, indicating significant interactions between factors and significant differences between groups, as noted in each item. The green line indicates the MD±SD of group C, and each scatter plot indicates a sampling unit. # indicates a significant interaction between liraglutide and kombucha. (**A**) Plasma triglyceride (TG) levels (n = 11 in the L-/K- and L+/K+ groups, and n = 10 in the L+/K- and L-/K+ groups). Two-Way ANOVA: L factor - F (1, 38) = 19.40, P < 0.0001; K factor - F (1, 38) = 2.487, P = 0.1231; Interaction - F (1, 38) = 0.3888, P = 0.5367. (**B**) log_10_-transformed plasma total cholesterol (TC) levels (n= 11 in L-/K+, L+/K+, L+/K-, and n = 9 in L-/K- groups). Two-Way ANOVA: L factor - F (1, 38) = 18.92, P < 0.0001; K factor - F (1, 38) = 0.0006980, P = 0.9791; Interaction - F (1, 38) = 0.1338, P = 0.7166. (**C**) Plasma high-density lipoprotein cholesterol (HDL-C) levels (n = 10 in L-/K-, and n = 11 in the other groups). Two-Way ANOVA: L factor - F (1, 39) = 5.330, P = 0.0263; K factor - F (1, 39) = 0.2944, P = 0.5905; Interaction - F (1, 39) = 0.8297, P = 0.3680. (**D**) log_10_-transformed plasma low-density lipoprotein cholesterol (LDL-C) levels (n= 6 in L-/K-, 8 in L-/K+, and 10 in L+/K- and L+/K+). Two-Way ANOVA: L factor - F (1, 30) = 115.6, P < 0.0001; K factor - F (1, 30) = 6.529, P = 0.0159; Interaction - F (1, 30) = 7.095, P = 0.0123. (**E**) Plasma very low-density lipoprotein cholesterol (VLDL-C) (n = 10 in L+/K- and L-/K+ groups, and n = 11 in L-/K- and L+/K+). Two-Way ANOVA: L factor - F (1, 38) = 19.21, P < 0.0001; K factor - F (1, 38) = 2.471, P = 0.1243; Interaction - F (1, 38) = 0.4009, P = 0.5304.

### 2.5. Liraglutide and kombucha improve liver histology in animals with diet-induced obesity

HFD was effective in increasing the liver area stained with Oil Red O (ORO), a dye that labels neutral lipids (Figure S4, A and B - C: 1.712 ± 0.338 vs. L-/K-: 28.324 ± 8.766%; P = 0.0024), suggesting the induction of hepatic steatosis in the animals. Additionally, the presence of vesicles, hepatocellular ballooning, and an inflammatory infiltrate was observed in liver samples from this group, as indicated by Hematoxylin and Eosin (H&E) staining, which was absent in samples from the control group (Figure S4, A). Furthermore, an increase in the liver area stained by Picrosirius Red (PSR) (Figure S5, A and C - C: 0.384 ± 0.041 vs. L-/K-: 1.616 ± 0.188%; P < 0.0001) was observed in the HFD-fed animals, concomitantly suggesting the induction of hepatic fibrosis. The factors liraglutide [Figure 4, A and B - L factor: F (1, 15) = 14.37, P = 0.0018] and kombucha [Figure 4, A and B - K factor: F (1, 15) = 4.952, P = 0.0418] showed an effect on the liver area stained by ORO, and their interaction was significant [Figure 4, A and B - Interaction: F (1, 15) = 4.623, P = 0.0483]. Thus, the combination of treatments was effective in reducing the accumulation of neutral lipids in the liver of obese animals (Figure 4, A and B - L+/K+: 8,534 ± 4,949 vs. L-/K-: 28,324 ± 8,766%; P = 0.0044), but its efficacy was not distinct from that of liraglutide (Figure 4, A and B - L+/K+: 8,534 ± 4,949 vs. L+/K-: 8,781 ± 7,654%; P > 0.9999 and Figure 4, A and B - L+/K-: 8,781 ± 7,654 vs. L-/K-: 28,324 ± 8,766%; P = 0.0030) or the effect promoted by kombucha (Figure 4, A and B - L+/K+: 8,534 ± 4,949 vs. L-/K+: 13,931 ± 6,096%; P = 0.6788 and Figure 4, A and B - L-/K+: 13,931 ± 6,096 vs. L- /K-: 28,324 ± 8,766%; P = 0.0279). Additionally, both the combination of liraglutide and kombucha and the isolated treatments resulted in the absence of hepatocellular ballooning and inflammatory infiltrates in the liver (Figure 5, A).

**Figure 4.**
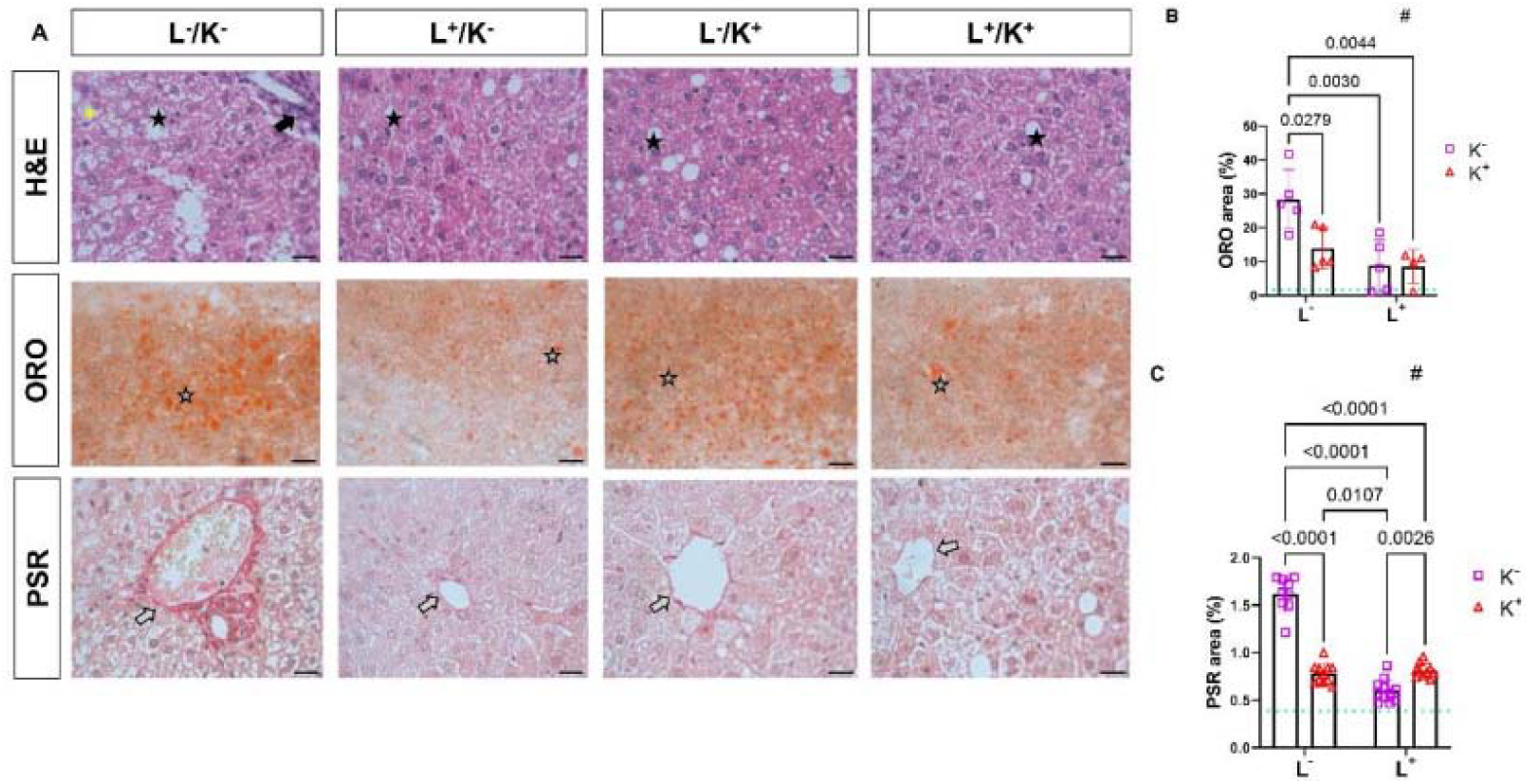
Hepatic histology. The results were obtained through a Two-Way ANOVA and are presented as mean ± standard deviation (SD), with a significance level of P ≤ 0.05. L factor = liraglutide factor; K factor = kombucha factor; interaction = interaction between factors. The results of the multiple comparison test are presented as P values, indicating significant interactions between factors and significant differences between groups, as noted in each item. The green line indicates the MD±SD of group C, and each scatter plot indicates a sampling unit. # indicates a significant interaction between liraglutide and kombucha. (**A**) Representative images of Hematoxylin and Eosin (H&E), Oil Red O (ORO), and Picrosirius Red (PSR) staining. (**B**) Hepatic area stained by ORO (n = 4 in L+/K+, and n = 5 in the other groups). Two-Way ANOVA: L factor - F (1, 15) = 14.37, P = 0.0018; K factor - F (1, 15) = 4.952, P = 0.0418; Interaction - F (1, 15) = 4.623, P = 0.0483. (C) Liver area stained by PSR transformed in log_10_ (n = 11 in all groups). Two-Way ANOVA: L factor - F (1, 37) = 155.4, P < 0.0001; K factor - F (1, 37) = 63.41, P < 0.0001; Interaction - F (1, 37) = 177.0, P < 0.0001. Magnification of 60× and scale bar of 17 µm in H&E, PSR and ORO. Black stars indicate the presence of vesicles, yellow asterisk indicates the presence of hepatocellular ballooning, black arrow indicates the presence of inflammatory infiltrate, gray stars indicate the accumulation of neutral lipids, and gray arrows indicate the presence of collagen fibers in the perivascular area.

**Figure 5.**
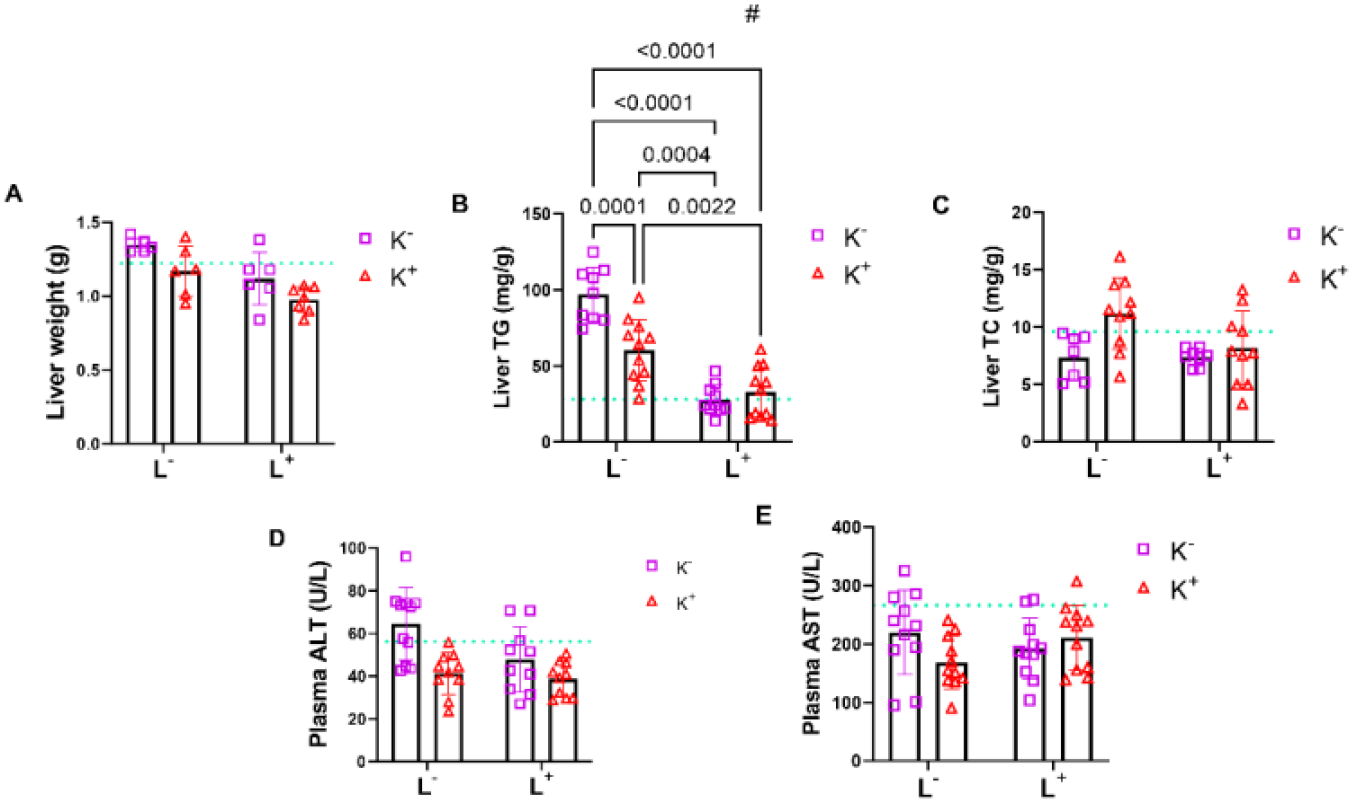
Hepatic parameters. The results were obtained through a Two-Way ANOVA and are presented as mean ± standard deviation (SD), with a significance level of P ≤ 0.05. L factor = liraglutide factor; K factor = kombucha factor; interaction = interaction between factors. The results of the multiple comparison test are presented as P values, indicating significant interactions between factors and significant differences between groups, as noted in each item. The green line indicates the MD±SD of group C, and each scatter plot indicates a sampling unit. # indicates a significant interaction between liraglutide and kombucha. (**A**) Liver weight (n = 7 in the L+/K+ group, and n = 6 in the other groups). Two-Way ANOVA: L factor - F (1, 21) = 16.04, P = 0.0006; K factor - F (1, 21) = 9.325, P = 0.0060; Interaction - F (1, 21) = 0.1155, P = 0.7374. (**B**) Hepatic TG content (n = 11 in L-/K+ and L+/K+ groups, n = 9 in L-/K-, and n = 10 in L+/K-). Two-Way ANOVA; L factor - F (1, 37) = 85.95, P < 0.0001; K factor - F (1, 37) = 9.123, P = 0.0046; Interaction - F (1, 37) = 15.91, P = 0.0003. (**C**) Hepatic TC content (n = 10 in L-/K+ and L+/K+ groups, n = 7 in L-/K-, and n = 8 in L+/K-). Two-Way ANOVA: L factor - F (1, 31) = 2.771, P = 0.1061; K factor - F (1, 31) = 6.798, P = 0.0139; Interaction - F (1, 31) = 2.925, P = 0.0972. (**D**) Plasma alanine aminotransferase (ALT) levels (n = 11 in the L-/K- group, and n = 10 in the other groups). Two-Way ANOVA: L factor - F (1, 37) = 5.519, P = 0.0243; K factor - F (1, 37) = 15.33, P = 0.0004; Interaction - F (1, 37) = 2.868, P = 0.0988. (**E**) Plasma aspartate aminotransferase (AST) levels (n = 11 in all groups). Two Way ANOVA: L factor - F (1, 40) = 0.1960, P = 0.6604; K factor - F (1, 40) = 0.8850, P = 0.3525; Interaction - F (1, 40) = 4.192, P = 0.0472.

Similarly, both the presence of liraglutide [Figure 4, A and C - L factor: F (1, 37) = 155.4, P < 0.0001] and kombucha [Figure 4, A and C - K factor: F (1, 37) = 63.41, P < 0.0001] affected the hepatic area stained by PSR, and the effects of these factors interacted significantly [Figure 4, A and C - Interaction: F (1, 37) = 177.0, P < 0.0001]. The combination of treatments reduced the area marked by collagen fibers in the liver of animals with steatosis (Figure 4, A and C - L+/K+: 0.811 ± 0.074 vs. L-/K-: 1.616 ± 0.188%; P < 0.0001), as well as liraglutide (Figure 4, A and C - L+/K-: 0.601 ± 0.120 vs. L-/K-: 1.616 ± 0.188%; P < 0.0001) and kombucha (Figure 4, A and C - L-/K+: 0.778 ± 0.104 vs. L-/K-: 1.616 ± 0.188%; P < 0.0001). The combination demonstrated lower efficacy compared to liraglutide (Figure 4, A and C - L+/K+: 0.811 ± 0.074 vs. L+/K-: 0.601 ± 0.120%; P = 0.0026), which also had a greater effect than kombucha (Figure 4, A and C - L+/K-: 0.601 ± 0.120 vs. L-/K+: 0.778 ± 0.104%; P = 0.0107).

### 2.6. Liraglutide and kombucha attenuate hepatic lipid accumulation in animals with MASLD

The HFD was effective at increasing liver weight (Figure S5, A- C: 1.222 ± 0.085 vs. L-/K-: 1.347 ± 0.046 g; P = 0.0100) and hepatic TG content (Figure S5, B - C: 28.064 ± 12.471 vs. L-/K-: 97.030 ± 17.866 mg/g; P < 0.0001), without altering hepatic TC content (Figure S5, C - C: 9.616 ± 2.826 vs. L-/K-: 7.348 ± 1.954 mg/g; P = 0.1292). Since HFD was not able to alter the plasma levels of the hepatic enzymes alanine aminotransferase (ALT) (Figure S5, D - C: 56,309 ± 9,470 vs. L-/K- : 64,523 ± 16,921 U/L; P = 0.3813) and aspartate aminotransferase (AST) (Figure S5, E - C: 266,265 ± 55,845 vs. L-/K-: 219,679 ± 72,230 U/L; P = 0.4117), the induction of a MASLD model in a transitional state between steatosis and MASH stages is suggested.

Both liraglutide [Figure 5, A - L factor: F (1, 21) = 16.04, P = 0.0006] and kombucha factors exerted an effect on liver weight [Figure 5, A - K factor: F (1, 21) = 9.325, P = 0.0060]; however, the interaction between both was not significant [Figure 5, A - Interaction: F (1, 21) = 0.1155, P = 0.7374].

Liraglutide [Figure 5, B - L factor: F (1, 37) = 85.95, P < 0.0001] and kombucha [Figure 5, B - K factor; F (1, 37) = 9.123, P = 0.0046] affected hepatic TG content, and the interaction between factors was significant [Figure 5, B - Interaction: F (1, 37) = 15.91, P = 0.0003]. Thus, the combination of treatments reduced hepatic TG content (Figure 5, B - L+/K+: 32,754 ± 16,754 vs. L-/K-: 97,030 ± 17,866 mg/g; P < 0.0001), and this effect was superior to that promoted by kombucha (Figure 5, B - L+/K+: 32,754 ± 16,754 vs. L-/K+: 60,374 ± 20,245 mg/g; P = 0.0022 and Figure 5, B - L-/K+: 60,374 ± 20,245 vs. L-/K-: 97,030 ± 17,866 mg/g; P = 0.0001), although it was not more effective than that of liraglutide (Figure 5, B - L+/K+: 32,754 ± 16,754 vs. L+/K-: 27,687 ± 9,667 mg/g; P = 0.8984 and Figure 5, B - L+/K-: 27,687 ± 9,667 vs. L-/K-: 97,030 ± 17,866 mg/g; P < 0.0001).

In contrast, kombucha-containing regimens [Figure 5, C - K factor: F (1, 31) = 6.798, P = 0.0139] showed a distinct effect from regimens without kombucha on hepatic TC content, which was not observed for liraglutide [Figure 5, C - L factor: F (1, 31) = 2.771, P = 0.1061]. The interaction between factors was not significant [Figure 5, C - Interaction: F (1, 31) = 2.925, P = 0.0972].

Similar to what was observed for liver weight, the effect of the factors liraglutide [Figure 5, D - L factor: F (1, 37) = 5.519, P = 0.0243] and kombucha [Figure 5, D - K factor: F (1, 37) = 15.33, P = 0.0004] on plasma ALT levels was not accompanied by a significant interaction [Figure 5, D - Interaction: F (1, 37) = 2.868, P = 0.0988].

Neither liraglutide [Figure 5, E - L factor: F (1, 40) = 0.1960, P = 0.6604] nor kombucha [Figure 5, E - K factor: F (1, 40) = 0.8850, P = 0.3525] had a significant effect on plasma AST levels, and although factor analysis indicated a significant interaction between such factors [Figure 5, E - Interaction: F (1, 40) = 4.192, P = 0.0472], the multiple comparison test did not detect a difference between any of the experimental groups.

### 2.7 HFD, Liraglutide, and Kombucha Do Not Modulate Hepatic Expression of autophagy-Related Proteins and LAL Activity in the Liver

The HFD did not affect the hepatic expression of the proteins involved in autophagy including Microtubule-Associated Protein 1A/1B-Light Chain 3 (LC3-II) (Figure S6, A and B - C: 0.128 ± 0.073 vs. L-/K-: 0.175 ± 0.067 A.U; P = 0.3214), Sequestosome-1 (p62) (Figure S6, A and C - C: 0.349 ± 0.164 vs. L-/K-: 0.313 ± 0.143 A.U; P = 0.7356), Autophagy Related 7 (Atg7) (Figure S6, A and D - C: 0.271 ± 0.171 vs. L-/K-: 0.383 ± 0.183 A.U; P = 0.3796), and LAL (Figure S6, A and E - C: 0.310 ± 0.218 vs. L-/K-: 0.193 ± 0.109 A.U; P = 0.3253) although an increase in the activity of this enzyme was noted with the consumption of HFD (Figure S6, F - C: 0.111 ± 0.016 vs. L-/K-: 0.144 ± 0.012 nmol/min/mg protein; P = 0.0083).

Neither the liraglutide factor [Figure 6, A and B - L factor: F (1, 19) = 0.002755, P = 0.9587], nor the kombucha factor [Figure 6, A and B - K factor: F (1, 19) = 1.433, P = 0.2460] affected the hepatic expression of LC3-II, nor was there a significant interaction between factors [Figure 6, A and B - Interaction - F (1, 19) = 1.681, P = 0.2103].

**Figure 6.**
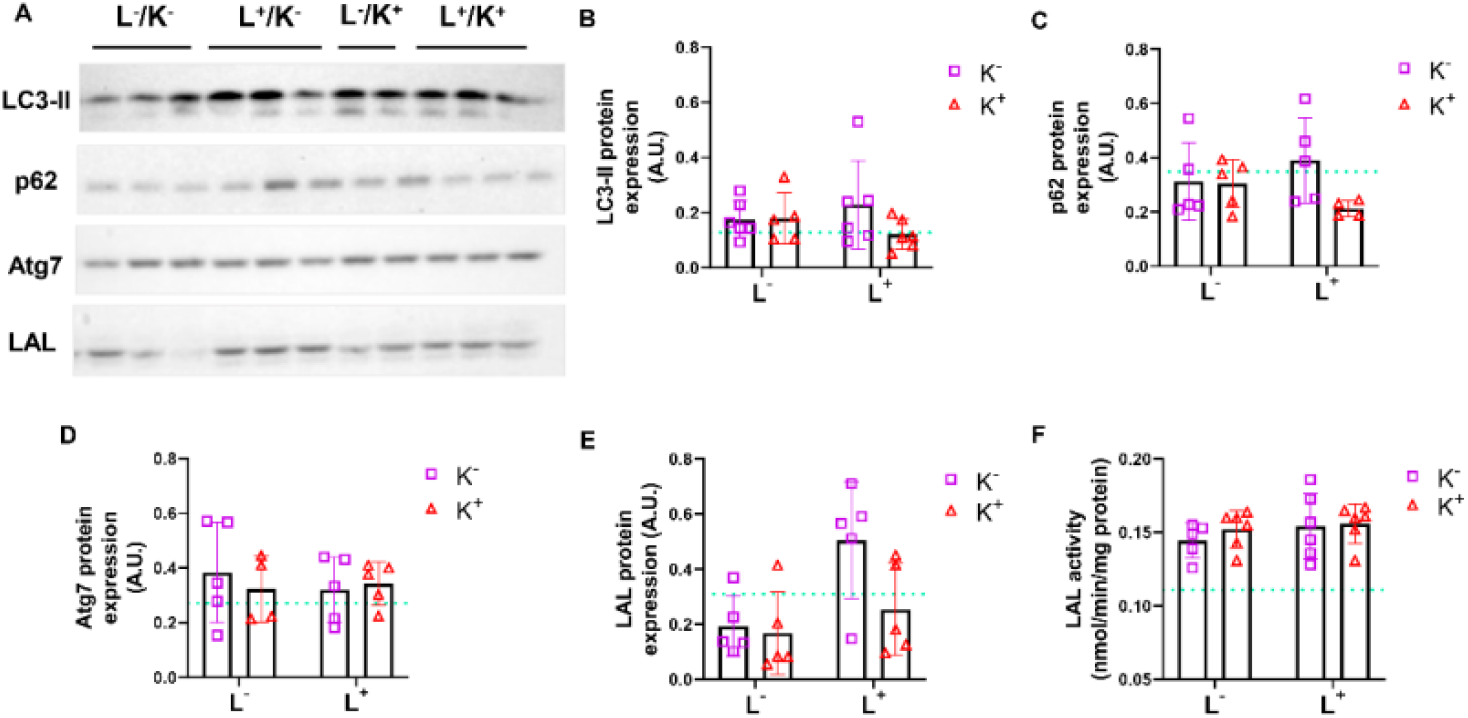
Hepatic autophagy-related proteins and lysosomal acid lipase (LAL) expression and activity. The results were obtained through a Two-Way ANOVA and are presented as mean ± standard deviation (SD), with a significance level of P ≤ 0.05. L factor = liraglutide factor; K factor = kombucha factor; interaction = interaction between factors. The results of the multiple comparison test are presented as P values, indicating significant interactions between factors and significant differences between groups, as noted in each item. The green line indicates the MD ± SD of group C, and each scatter plot indicates a sampling unit. # indicates a significant interaction between liraglutide and kombucha. (**A**) Representative image of the hepatic expression of Microtubule-Associated Protein 1A/1B-Light Chain 3 (LC3-II), Sequestosome-1 (p62), Autophagy Related 7 (Atg7), and LAL. (**B**) Hepatic expression of LC3-II, obtained by Western blotting (n = 5 in the L-/K+ group, and n = 6 in the other groups). Two-Way ANOVA: L factor - F (1, 19) = 0.002755, P = 0.9587; K factor - F (1, 19) = 1.433, P = 0.2460; Interaction - F (1, 19) = 1.681, P = 0.2103. (**C**) Hepatic expression of p62, obtained by Western blotting (n = 4 in the L+/K+ group, and n = 5 in the other groups). Two-Way ANOVA: L factor - F (1, 15) = 0.01714, P = 0.8976; K factor - F (1, 15) = 2.825, P = 0.1135; Interaction - F (1, 15) = 2.376, P = 0.1441. (**D**) Hepatic expression of Atg7, obtained by Western blotting (n = 4 in the L-/K+ group, and n = 5 in the other groups). Two-Way ANOVA: L factor - F (1, 15) = 0.1246, P = 0.7290: K factor - F (1, 15) = 0.09354, P = 0.7639; Interaction - F (1, 15) = 0.4621, P = 0.5070. (**E**) Hepatic expression of LAL, obtained by Western blotting (n = 5 in all groups). Two-Way ANOVA: L factor - F (1, 16) = 7.382, P = 0.0152; K factor - F (1, 16) = 3.559, P = 0.0775; Interaction - F (1, 16) = 2.388, P = 0.1418. (F) LAL activity in the liver, obtained from fluorescence assay (n = 5 in the L-/K- group, and n = 6 in the other groups). Two-Way ANOVA: L factor - F (1, 19) = 1.060, P = 0.3162; K factor - F (1, 19) = 0.5281, P = 0.4763; Interaction - F (1, 19) = 0.2017, P = 0.6584. Two independent membranes were used for statistical analysis (n total 4–6 per group). Image of membranes representing all groups (C, L-/K-, L+/K-, L-/K+ and L+/K+), cropped for representative purposes in this figure. The same samples from the L-/K- group were used in both this figure and Figure S6, A.

The same was observed for the proteins p62 [Figure 6, A and C - L factor: F (1, 15) = 0.01714, P = 0.8976; K factor: F (1, 15) = 2.825, P = 0.1135; Interaction: F (1, 15) = 2.376, P = 0.1441] and Atg7 [Figure 6, A and D - L factor: F (1, 15) = 0.1246, P = 0.7290; K factor: F (1, 15) = 0.09354, P = 0.7639; Interaction: F (1, 15) = 0.4621, P = 0.5070]. Liraglutide factor had an effect on LAL expression, unlike kombucha factor, and there was no significant interaction between both [Figure 6, A and E - L factor: F (1, 16) = 7.382, P = 0.0152; K factor: F (1, 16) = 3.559, P = 0.0775; Interaction: F (1, 16) = 2.388, P = 0.1418]. The factors liraglutide [Figure 6, F - L factor: F (1, 19) = 1.060, P = 0.3162] and kombucha [Figure 6, F - K factor: F (1, 19) = 0.5281, P = 0.4763] had no effect on LAL activity in the liver, and there was no significant interaction between the two [Figure 6, F - Interaction: F (1, 19) = 0.2017, P = 0.6584].

## 3. Discussion

The effects of liraglutide on body weight have already been described in the literature, corroborating the data observed here. Indeed, C57BL/6 mice fed HFD for 13 weeks and treated with the same daily dose of liraglutide and for the same duration as in the present study also experienced significant body weight loss [15]. Moreover, there is evidence that kombucha has an effect that improves liver parameters in MASLD. However, these improvements were not accompanied by a reduction in body weight [12,14]. These findings are consistent with the data obtained in the present study, which shows the notable efficacy of the GLP-1 agonist on the body dynamics of the animals, without an enhanced effect in combination with the probiotic drink.

The effect of kombucha on increasing satiety in mice has been previously reported in the literature [16]. The significant interaction between liraglutide and kombucha observed in the dietary parameters studied here suggests a potential synergistic effect between these treatments. However, the effect promoted by kombucha was not more effective than that of regimens containing only liraglutide, demonstrating the superior efficacy of liraglutide in reducing food intake. Therefore, in addition to binding to gastric receptors, the drug acts on cortical areas involved in the control of food intake, promoting this effect [17-19].

The treatment effects observed here are consistent with data described in the literature, indicating effects of liraglutide and kombucha on glycemic homeostasis. For example, after 9 days of treatment, kombucha reduced fasting blood glucose and the AUC of the glucose tolerance test performed in C57BL/6 mice with HFD-induced MASLD, indicating that this beverage influences glucose tolerance and basal blood glucose control, contributing to the improvement of the disease [14]. Additionally, studies evaluating the effect of liraglutide on glycemic parameters in experimental MASH models reported that after 10 weeks of treatment, liraglutide reduced fasting blood glucose in C57BL/6J mice with HFD-induced T2DM and MASH and that after 4 weeks of treatment with liraglutide, obese db/db mice, with diabetes and steatosis, also displayed improvements in these same parameters [20, 21].

Regarding fasting blood glucose, we observed a superior effect of liraglutide compared to kombucha. The impact of liraglutide and kombucha on glucose tolerance, indicated by factor analysis and accompanied by a non-significant interaction, may suggest that these treatments act independently on this parameter, with no additional effect from the combined administration.

The effects of liraglutide on the lipid profile have been previously reported in clinical and preclinical trials. One study reported a reduction in plasma TG levels after 6 months of drug administration in a cohort of obese patients with T2DM [22]. Other reports of liraglutide’s impact on circulating TG levels have also been described in the literature, although its effect on the levels of other lipids in individuals with T2DM has not been reported [23]. On the other hand, in murine models of T2DM and other metabolic diseases, liraglutide had an impact on other types of circulating lipids. For example, when administered for 4 weeks to Sprague–Dawley rats with steatosis induced by a diet rich in lipids and cholesterol, there was a reduction in serum levels not only of TG, but also of TC [24].

The lipid-lowering effects attributed to kombucha and its influence on the lipid profile in preclinical studies has yielded controversial results [25]. On the one hand, when administered to Wistar rats with dyslipidemia at a dose of 1.8 mL twice daily for 14 days, kombucha did not reduce plasma levels of HDL-C and LDL-C [26]. On the other hand, treating rats of the same lineage with the beverage for 16 weeks at a dose of 5 mL/kg body weight/day resulted in an attenuation of circulating levels of TC, LDL-C, VLDL-C, and TAG [27]. These data may suggest that the effects of kombucha on the lipid profile require a longer administration period, at least two weeks, to manifest. In this study, we did not observe an additional effect resulting from the combination of treatments on any of the lipid parameters evaluated.

Improvement in steatosis promoted by liraglutide after various administration regimens and at different doses has already been reported in other preclinical studies, based on liver staining with ORO [28-30]. The results observed here agree with those reported in these studies. Kombucha also showed significant effects on improving steatosis from this histological technique, indicating the impact of both treatments on reducing the accumulation of neutral lipids in the liver [12,14]. In the present study, no difference was found between the effects promoted by liraglutide and those promoted by kombucha, and the combination of both did not offer any additional benefit.

The effects of liraglutide on hepatic fibrosis observed here align with some clinical trials, which indicate that this drug is beneficial for patients with T2DM and MASH [31,32]. This drug also improved this parameter in animal models of MASLD [33]. Although some authors also indicate an improvement in this parameter when treated with kombucha, we found that liraglutide proved to be more effective than the combination of treatments in reducing the hepatic area stained by PSR, indicating that the combination of supplementation with the GLP-1 analogue may have exerted an antagonistic action [12,14].

In addition to the effects observed in the liver area stained by ORO, the effects of liraglutide and kombucha on liver weight and hepatic TG content indicate that both treatments are effective in reducing hepatic steatosis. Liraglutide proved more effective than kombucha in reducing hepatic TG, and the combination of both did not provide an additional effect beyond that observed with liraglutide, highlighting the drug’s effect on attenuating the early stage of MASLD. None of the treatments altered the hepatic content of TC, ALT, and AST, possibly because the induced model was at an early stage of MASLD, characterized by hepatic steatosis without significant hepatocellular injury, as HFD was also unable to alter these parameters.

The literature suggests that LAL plays a crucial role in intracellular lipid catabolism within lysosomes, regulating the hydrolysis of cholesterol esters and TG, thereby producing cholesterol and fatty acids; though, some results are conflicting [34]. According to some studies, plasma LAL activity is inversely associated with the progression of MASLD and correlated with hepatic LAL activity [35- 37]. Moreover, specific knockout of LAL in the liver resulted in increased hepatic lipid accumulation [38]. However, hepatic LAL overexpression was unable to attenuate steatosis in the livers of mice fed a Western diet or upregulate the hepatic expression of inflammatory genes [39]. Thus, the results obtained in this study indicate that HFD increased LAL activity and did not significantly alter the hepatic expression of autophagy-related proteins (e.g., LC3-II, p62, and Atg7).

Although liraglutide impacted LAL expression, no interaction was observed between the factors, and none of the other lipophagy markers assessed were modulated by the interventions. It is important to consider that the fasting period used before euthanasia may have limited the detection of effects on lipophagy, since a 12–18 hour food restriction period is reported in autophagy induction protocols [28,40,41]. Thus, the data suggest that under the conditions tested, HFD and the treatments had no impact on the lipophagic pathway in the liver. The importance of studying the effects of different fasting periods on the induction of lipophagy in murine models, as well as the effects promoted by liraglutide, kombucha, and their combination on hepatic lipophagy, is highlighted, towards the goal of elucidating the potential mechanisms involved in the improvement of MASLD observed with these agents.

## 4. Materials and Methods

### 4.1. Animals

Seven-week-old male C57BL/6 mice were provided by the animal facility of the University of São Paulo School of Medicine (FMUSP). All animals were housed in specific isolated cages in a room with controlled temperature of 25 ± 2 °C and a 12:12 h light-dark cycle (6:00 am to 6:00 pm), and had ad libitum access to food and water. After 1 week of acclimatization, the animals were randomized using the Random Sequence virtual resource (https://www.random.org/sequences/) and divided into two groups: (1) animals fed a control diet (C), containing 19% protein, 56% carbohydrates, 3.5% lipids, 5% cellulose, and 4.5% vitamins and minerals, providing 3.20 kcal/g (CR-1, Nuvilab, Colombo, Paraná, Brazil) and (2) animals fed a pelleted HFD, containing 15.8% protein, 27% carbohydrates, 57.2% lipids, 5% cellulose, and 4.5% vitamins and minerals, providing 5.35 kcal/g (PragSoluções Biociências, Jaú, São Paulo, Brazil). These groups of mice received their respective diets consecutively for 10 weeks. At the end of this period, the HFD-fed animals were randomized and divided into four groups: (1) animals receiving HFD (L-/K-), (2) animals fed HFD and treated with liraglutide (L+/K-), (3) animals fed HFD and treated with kombucha (L-/K+), and (4) animals fed HFD and treated simultaneously with liraglutide and kombucha (L+/K+). These interventions lasted for 15 consecutive days. The use of animals in this study was supported by approval from the Ethics Committee for Animal Use of the Institute of Biomedical Sciences of the University of São Paulo (CEUA/ICB), under protocol number 3819110222, approved on April 26, 2022.

### 4.2. Liraglutide Treatment

The animals received liraglutide at a dose of 200 μg/kg body weight, administered twice daily (8:00 am and 3:00 pm) via subcutaneous injection, for 15 consecutive days [42,43]. The drug was commercially available under the name Victoza (Novo Nordisk, Bagsværd, Denmark). The animals’ body weight and food intake were monitored daily during treatment with liraglutide.

### 4.3. Kombucha Treatment

Six grams of green tea (Camellia sinensis) were added to 1000 mL of boiling distilled water, allowed to steep for 5 min, and then strained. Sixty grams of sucrose were added and stirred to dissolve it in the green tea. After reaching room temperature, the tea was poured into a previously sterilized glass container and heated to 120 °C for 20 minutes. Finally, the freshly brewed tea was inoculated with a symbiotic kombucha culture containing 100 mL of previously fermented tea. The container was covered with gauze to prevent external insect contamination and allow aerobic fermentation. Fermentation (25 ± 3 °C) continued for 10 days, until the pH reached 3.1–3.2 and the Brix level was 4–5%. The kombucha colony was obtained commercially from Kombucha Kamp (Gardena, CA, USA). The kombucha was collected and centrifuged at 7,000 × g for 20 min, and the supernatant was passed through a 0.22 μm syringe filter (Millipore, Burlington, MA, USA). The daily dose administered to each mouse corresponded to 0.2 mL containing 10^7^–10^8^ microorganisms/mL of kombucha, which was determined spectrophotometrically by measuring the absorbance at 600 nm. The treatment was administered once a day (9:00 am) for 15 consecutive days. The body weight and food intake of the animals were monitored daily during the kombucha treatment [14,44].

### 4.4. oGTT

At the end of the treatment period, and after a 6-hour fast, the animals, while awake, had a drop of blood collected after sectioning their tails to measure their basal blood glucose levels. Blood glucose was determined using test strips (Roche, Mannheim, Germany) and a strip reading device (Accu-chek Active; Roche). The animals then received, orally (gavage), a solution containing 25% glucose at a dose of 75 mg/100 g body weight [45]. In addition to basal blood glucose, new blood collections for plasma glucose measurements were performed at 5, 10, 15, 30, 45, 60, and 90 minutes after glucose administration. The aliquots for determining blood glucose levels were measured as described previously [46,47].

### 4.5. Body composition

On the day after the last day of treatment, the animals underwent magnetic resonance imaging (MRI) body composition analysis using a Minispec LF50 (Bruker, Billerica, MA, USA). The parameters analyzed were lean and fat mass.

### 4.6. Euthanasia

On the day following the last day of treatment (20 weeks of age), and after a 6-hour fast, the animals were anesthetized with isoflurane. The depth of anesthesia was assessed by monitoring motor reflexes. After ensuring sedation, rapid decapitation was performed [48]. After weighing the liver, liver tissue samples were collected, quickly placed in a liquid nitrogen (N2) bath, and subsequently stored at –80 °C. Some liver samples were also fixed in a 10% formaldehyde solution for 24 hours, as previously described, or embedded in Tissue-Tek (Thermo Scientific, Waltham, MA, USA) [49]. Blood samples were also collected, quickly placed on ice and stored at “80 °C for later use.

### 4.7. Plasma Lipids

After euthanasia, the animals’ blood was collected and centrifuged (4 °C, 4,000 × g, 30 minutes) to separate the plasma. Subsequently, plasma TG, TC, and HDL-C concentrations were determined by enzymatic colorimetry, using commercial kits (TG: Cat. No. 87; TC: Cat. No. 76; HDL-C: Cat. No. 145, LabTest, Belo Horizonte, Minas Gerais, Brazil) according to the manufacturer’s instructions. Plasma VLDL-C and LDL-C concentrations were determined as previously described in the literature [50].

### 4.8. H&E, PSR e ORO

For H&E and PSR staining, liver samples were fixed in 10% formaldehyde solution for 24h in individual cassettes after euthanasia [49]. Subsequently, the fixed samples were kept overnight in 70% alcohol, dehydrated through a series of baths in 95% alcohol, 100% alcohol, and xylene, and embedded in paraffin at 60 °C. A microtome (Zeiss, Jena, Germany) was used to cut the samples into 5-μm sections.

The sections were then stained with H&E to analyze liver morphology and with PSR to identify collagen fibers, as previously described [51]. The H&E-stained samples underwent qualitative analysis for the presence of vesicles, hepatocellular ballooning, and inflammatory infiltrates.

The samples intended for staining with ORO, a fat-soluble dye that marks neutral lipids, were embedded in Tissue-Tek, placed in containers filled with isopropanol, and immediately immersed in a liquid N2 bath [52,53]. Twelve μm sections were prepared using a cryostat (Microm H560; Thermo Scientific).

For H&E staining, twenty images were analyzed, and for PSR and ORO ten images were analyzed with a 20× objective (three distinct sections) of each animal. H&E, PSR and ORO images were obtained using a Nikon Eclipse Ti-U microscope coupled to a Nikon DS-R1 digital camera and NIS Elements BR 3.1 software

The abundance of collagen deposition and lipid accumulation in the liver was presented as the mean percentage for each animal within each group. The identification of the PSR-stained and the ORO-stained areas were performed using the ImageJ v1.8.0 software (National Institutes of Health-NIH, Bethesda, MD, USA). First, the image was opened in the program and converted to 8-bit grayscale (Image ⟶ type ⟶ 8 bits). Then, the following sequence was performed: Image ⟶ Adjust ⟶ Threshold; Analyze ⟶ Define measurements. The results were obtained through the commands “Analyze” ⟶ “Measure”. Area values, which correspond to the amount of collagen or lipid accumulation, were recorded [52]. For representative purposes, images of the H&E, PSR, and ORO-stained samples were also obtained with a 60× objective.

### 4.9. Liver TG and CT Content

After euthanasia, liver samples stored at –80 °C underwent lipid extraction, as previously described [54]. After extraction, total TG and CT were quantified through enzymatic colorimetry using specific commercial kits (TG: Cat. No. 87; CT: Cat. No. 76, LabTest, Belo Horizonte, Minas Gerais, Brazil) according to the manufacturer’s instructions.

### 4.10. Plasma ALT and AST

To investigate hepatocellular damage, the concentrations of the liver enzymes ALT and AST were measured in centrifuged blood (4 °C, 4,000 × g, 30 min) using the Kinetic-UV method, using commercial kits, according to the manufacturer’s instructions (ALT: Cat. No. 108 and AST: Cat. No. 109, LabTest, Belo Horizonte, Minas Gerais, Brazil).

### 4.11. Hepatic LAL Activity

Liver samples were homogenized, and hepatic LAL activity was quantified by fluorescence, according to the manufacturer’s instructions for the kit used (Cat. No. 700640; Cayman Chemical, Ann Arbor, MI, USA).

### 4.12 .Western blotting

Liver samples were homogenized in a lysis buffer containing phosphatase/protease inhibitors. The homogenate was placed on ice for 30 minutes and then centrifuged at 12,000 rpm for 40 minutes at 4 °C. The supernatant was collected, and the total protein concentration was measured using the Quick Start protein assay kit (Cat. No. 500-0205; Bio-Rad, Hercules, CA, USA). Electrophoresis was performed on a 10 or 20% polyacrylamide SDS-PAGE gel (83 × 73 × 15 mm) and subsequently transferred to a nitrocellulose membrane. The membranes were blocked overnight in a 5% powdered milk solution (Molico, Nestlé, São Paulo, Brazil) to avoid nonspecific protein binding. The membranes were incubated overnight at 4 °C with the primary antibodies p62 (Cat. No. 23214, Cell Signaling, Danvers, MA, USA, 1:1000), LC3A/B (Cat. No. 12741, Cell Signaling, Danvers, MA, USA, 1:1000), Atg7 (Cat. No. 8558, Cell Signaling, Danvers, MA, USA, 1:1000), and LAL (Cat. No. 12956-1- AP, Proteintech, Rosemont, IL, USA, 1:500). Then, the membranes were washed and incubated with a peroxidase-conjugated secondary antibody for 75 min at room temperature. Two independent membranes were obtained and analyzed for each protein analyzed, both containing samples from all experimental groups. Densitometric analysis of Western blot bands was performed using ImageJ v1.8.0 software (NIH). All samples were normalized by staining nitrocellulose membranes with Ponceau S.

### 4.13. Data Analysis

Data were analyzed using GraphPad Prism version 9.0 software (GraphPad Software, La Jolla, CA, USA). Initially, data were assessed for normal distribution using the Shapiro–Wilk test and for homoscedasticity using the Brown–Forsythe test. Subsequently, the data were subjected to a two-way ANOVA, followed by Tukey’s post hoc test. When the assumptions of normality and homoscedasticity were not met, the data underwent a base-10 logarithm (log_10_) transformation.

The data used to characterize the model (supplementary material) were tested for normal distribution using the Shapiro–Wilk test and for homoscedasticity using the Brown–Forsythe test, and were subsequently subjected to the unpaired Student’s t-test or nonparametric version. The significance level was set at 5% (P ≤ 0.05).

## 5. Conclusions

In summary, the results of this study corroborate the literature findings by demonstrating the superior efficacy of liraglutide in reducing body weight, food intake, and glycemic parameters in a murine model of MASLD. Despite kombucha’s beneficial effects on glycemic homeostasis and hepatic steatosis, its combination with the drug did not result in a significant additional effect. Although an effect on LAL expression was observed with the presence of liraglutide, the treatments did not modulate markers of the lipophagic pathway or LAL activity under the conditions tested, suggesting that their mechanisms of action likely involve other metabolic pathways. Therefore, it is concluded that liraglutide remains the most potent intervention, without its efficacy being optimized by kombucha supplementation.

## Author Contributions

J.S.A. and C.R.O.C. conceived and designed the study. J.S.A. performed the experiments, analyzed the data, and wrote the manuscript. S.A., T.Y.O., B.L.C.F., R.S.O.G., F.N.C. and A.C.M.K. contributed to data collection. C.R.O.C. supervised the project and revised the manuscript. All authors read and approved the final version.

## Funding

This research was funded by the National Council for Scientific and Technological Development (CNPq) (grant number 140217/2022-3) and São Paulo Research Foundation (FAPESP) (grant numbers 2016/25129-4; 2017/18972-0 and 2021/10469-2).

## Institutional Review Board Statement

The animal study protocol was approved by the Ethics Committee for Animal Use of the Institute of Biomedical Sciences of the University of São Paulo (CEUA/ICB) (protocol number 3819110222, approved on April 26, 2022).

## Acknowledgments

We thank the National Council for Scientific and Technological Development (CNPq) and São Paulo Research Foundation (FAPESP) for financial support. We also thank Cleide Rosana Duarte Prisco, department statistician, for her assistance with data analysis, and Adilson da Silva Alves, specialist at the Cellular Neurobiology Laboratory, for his assistance with Oil Red O staining. We would also like to thank Gustavo Cândido e Cruz, M.Sc.; Beatriz Garcia de Carvalho, M.Sc.; Sandro Leão Matos, Ph.D.; and Andressa Godoy Amaral, Ph.D.,for their initial support during the experimental phase.

## Conflicts of Interest

The authors declare that they have no conflicts of interest.

## Abbreviations

MASLD: Metabolic dysfunction–associated steatotic liver disease
LDL-C: Low-density lipoprotein cholesterol
HFD: High-fat diet
MASH: Metabolic dysfunction–associated steatohepatitis
LAL: Lysosomal acid lipase
GLP-1: Glucagon-like peptide-1
FDA: Food and Drug Administration
T2DM: Type 2 diabetes mellitus
AUC: Area under the curve
log_10_: Base-10 logarithm
oGTT: Oral glucose tolerance test
TG: Triglycerides
TC: Total cholesterol
HDL-C: High-density lipoprotein cholesterol
VLDL-C: Very low-density lipoprotein cholesterol
ORO: Oil Red O
H&E: Hematoxylin and Eosin
PSR: Picrosirius Red
ALT: Alanine aminotransferase
AST: Aspartate aminotransferase
LC3-II: Microtubule-Associated Protein 1A/1B-Light Chain 3
p62: Sequestosome-1
Atg7: Autophagy Related 7

## Notes

### Competing Interest Statement

The authors have declared no competing interest.

### Summary of Updates

Aiming for better preprint quality and to promote open science, two authors were added and the acknowledgments section was updated.

## References

1. Moreira, R.O.; Zagury, R.L.; Nogueira, I.C.R.; Nunes, V.S.; Salles, J.E.N.; Oliveira, C.P.M.S.; Coutinho, W.F.; de Faria, E.C.; de Faria, V.C.R.; de Godoy, J.R.P.; et al. Brazilian evidence-based guideline for screening, diagnosis, treatment, and follow-up of metabolic dysfunction-associated steatotic liver disease (MASLD) in adult individuals with overweight or obesity: a joint position statement from the brazilian society of endocrinology and metabolism (sbem), brazilian society of hepatology (sbh), and brazilian association for the study of obesity and metabolic syndrome (abeso). Arch. Endocrinol. Metab. 2023, 67, 865–882, 10.20945/2359-4292-2023-0123.

2. Sangro, P.; Dettori, S.; Cossu, E.; Pujia, A.; Meroni, M. Metabolic dysfunction–associated fatty liver disease (MAFLD): an update of the recent advances in pharmacological treatment. J. Physiol. Biochem. 2023, 79, 869–886, 10.1007/s13105-023-00954-4.

3. Yanai, H.; Adachi, H.; Hakoshima, M.; Katsuyama, H. Metabolic-Dysfunction-Associated Steatotic Liver Disease—Its Pathophysiology, Association with Atherosclerosis and Cardiovascular Disease, and Treatments. Int. J. Mol. Sci. 2023, 24, 15473, 10.3390/ijms242015473.

4. Miao, L.; Chen, Y.; Qiu, Y.; Wang, J.; Li, Y. Current status and future trends of the global burden of MASLD. Trends Endocrinol. Metab. 2024, 35, 130–140, 10.1016/j.tem.2024.02.007.

5. Gluchowski, N.L.; Becuwe, M.; Walther, T.C.; Farese, R.V. Lipid droplets and liver disease: from basic biology to clinical implications. Nat. Rev. Gastroenterol. Hepatol. 2017, 14, 343–355, 10.1038/nrgastro.2017.32.

6. Jakubek, P.; Kejík, Z.; Kaplánek, R.; Veselá, E.; Abramenko, N.; Hromádka, R.; Masarík, M. Autophagy alterations in obesity, type 2 diabetes, and metabolic dysfunction-associated steatotic liver disease: the evidence from human studies. Intern. Emerg. Med. 2024, 10.1007/s11739-024-03700-w.

7. Zechner, R.; Madeo, F.; Kratky, D. Cytosolic lipolysis and lipophagy: two sides of the same coin. Nat. Rev. Mol. Cell Biol. 2017, 18, 671–684, 10.1038/nrm.2017.76.

8. Zhang, S.; Peng, X.; Yang, S.; Li, X.; Huang, M.; Wei, S.; Liu, J.; He, G.; Zheng, H.; Yang, L.; et al. The regulation, function, and role of lipophagy, a form of selective autophagy, in metabolic disorders. Cell Death Dis. 2022, 13, 132, 10.1038/s41419-022-04593-3.

9. Dikic, I.; Elazar, Z. Mechanism and medical implications of mammalian autophagy. Nat. Rev. Mol. Cell Biol. 2018, 19, 349–364, 10.1038/s41580-018-0003-4.

10. Attia, S.L.; Softic, S.; Mouzaki, M. Evolving Role for Pharmacotherapy in NAFLD/NASH. Clin. Transl. Sci. 2020, 14, 11–19, 10.1111/cts.12839.

11. Hyun, J.; Lee, Y.; Wang, S.; Kim, J.; Kim, J.; Cha, J.; Seo, Y.; Jung, Y. Kombucha tea prevents obese mice from developing hepatic steatosis and liver damage. Food Sci. Biotechnol. 2016, 25, 861–866, 10.1007/s10068-016-0142-3.

12. Lee, C.; Kim, J.; Wang, S.; Sung, S.; Kim, N.; Lee, H.H.; Seo, H.; Jung, Y. Hepatoprotective Effect of Kombucha Tea in Rodent Model of Nonalcoholic Fatty Liver Disease/Nonalcoholic Steatohepatitis. Int. J. Mol. Sci. 2019, 20, 2369, 10.3390/ijms20092369.

13. Zhang, F.; Wang, H.; Li, Y.; Wang, L.; Liu, X.; Li, Y.; Hou, Y.; Wang, X.; Wang, Y.; Wang, L.; et al. Probiotic Mixture Ameliorates a Diet-Induced MASLD/MASH Murine Model through the Regulation of Hepatic Lipid Metabolism and the Gut Microbiome. J. Agric. Food Chem. 2024, 72, 8613–8627, 10.1021/acs.jafc.3c08910.

14. Moreira, G.V.; Azevedo, F.F.; Ribeiro, L.M.; Santos, A.; Guadagnini, D.; Gama, P.; Carvalho, C.R.O. Kombucha tea improves glucose tolerance and reduces hepatic steatosis in obese mice. Biomed. Pharmacother. 2022, 155, 113660, 10.1016/j.biopha.2022.113660.

15. Bugáňová, M.; Hrenák, J.; Ochodnicka-Mackovicova, K.; Krajcirovicova, K.; Pesta, M.; Křen, V.; Cahová, M. The effects of liraglutide in mice with diet-induced obesity studied by metabolomics. J. Endocrinol. 2017, 233, 93–103, 10.1530/JOE-16-0478.

16. Urrutia, M.A.D.; da Silveira, T.R.; de Mello, A.S.; da Silva, J.; da Rosa, L.C.; de Oliveira, A.; de Oliveira, M.F.V.; de Souza, C.G.; da Silva, M.M.; de Souza, R.L.R. Effects of supplementation with kombucha and green banana flour on Wistar rats fed with a cafeteria diet. Heliyon 2021, 7, e07081, 10.1016/j.heliyon.2021.e07081.

17. Alruwaili, H.; Dehestani, B.; le Roux, C.W. Clinical Impact of Liraglutide as a Treatment of Obesity. Clin. Pharmacol. 2021, 13, 53–60, 10.2147/CPAA.S276085.

18. Farr, O.M.; Tsoukas, M.A.; Triantafyllou, G.; Dincer, F.; Filippaios, A.; Ko, B.J.; Mantzoros, C.S. GLP-1 receptors exist in the parietal cortex, hypothalamus and medulla of human brains and the GLP-1 analogue liraglutide alters brain activity related to highly desirable food cues in individuals with diabetes: a crossover, randomised, placebo-controlled trial. Diabetologia 2016, 59, 954–965, 10.1007/s00125-016-3874-y.

19. Larsen, P.J.; Tang-Christensen, M.; Holst, J.J.; Ørskov, C. Distribution of glucagon-like peptide-1 and other preproglucagon-derived peptides in the rat hypothalamus and brainstem. Neuroscience 1997, 77, 257–270, 10.1016/s0306-4522(96)00434-4.

20. Yang, P.; Zhang, L.; Zhang, Y.; He, J. Liraglutide ameliorates nonalcoholic fatty liver disease in diabetic mice via the IRS2/PI3K/Akt signaling pathway. Diabetes Metab. Syndr. Obes. 2019, 12, 1013–1021, 10.2147/DMSO.S206867.

21. Liu, Q.; Li, Y.; Liu, J.; Chen, Y.; Wu, X.; Wu, M.; Zhang, Y.; Li, X.; Cao, G. Liraglutide modulates gut microbiome and attenuates nonalcoholic fatty liver in db/db mice. Life Sci. 2020, 261, 118457, 10.1016/j.lfs.2020.118457.

22. Varanasi, A.; Patel, P.; Makdissi, A.; Dhindsa, S.; Chaudhuri, A.; Dandona, P. Clinical Use of Liraglutide in Type 2 Diabetes and its Effects on Cardiovascular Risk Factors. Endocr. Pract. 2012, 18, 140–145, 10.4158/EP11169.OR.

23. Courrèges, J.-P.; Vilsbøll, T.; Zdravkovic, M.; Le-Thi, T.; Krarup, T.; Schmitz, O.; Verhoeven, R.; Bugáňová, I.; Madsbad, S. Beneficial effects of once-daily liraglutide, a human glucagon-like peptide-1 analogue, on cardiovascular risk biomarkers in patients with Type 2 diabetes. Diabet. Med. 2008, 25, 1129–1131, 10.1111/j.1464-5491.2008.02484.x.

24. Gao, H.; Zeng, Z.; Zhang, H.; Zhou, X.; Guan, L.; Deng, W.; Xu, L. The Glucagon-Like Peptide-1 Analogue Liraglutide Inhibits Oxidative Stress and Inflammatory Response in the Liver of Rats with Diet-Induced Non-alcoholic Fatty Liver Disease. Biol. Pharm. Bull. 2015, 38, 694–702, 10.1248/bpb.b14-00505.

25. Abaci, N.; Senol Deniz, F.S.; Orhan, I.E. Kombucha – An ancient fermented beverage with desired bioactivities: a narrowed review. Food Chem. X 2022, 14, 100302, 10.1016/j.fochx.2022.100302.

26. Darmawan, A.E.; Kusuma, H.S.; Putri, D.K.; Mahfud, M. Effect of rosella-based kombucha tea on the lipid profile on hyperlipidemic rats (Rattus norvegicus). Niche J. Trop. Biol. 2018, 1, 42–47, 10.14710/niche.1.2.42-47.

27. Bellassoued, K.; Ghrab, F.; Makni-Ayadi, F.; Van Pelt, J.; Elfeki, A.; Ammar, E. Protective effect of kombucha on rats fed a hypercholesterolemic diet is mediated by its antioxidant activity. Pharm. Biol. 2015, 53, 1699–1709, 10.3109/13880209.2014.1001408.

28. Fang, Y.; Chen, H.; Wang, C.; Liang, L. Liraglutide Alleviates Hepatic Steatosis by Activating the TFEB-Regulated Autophagy-Lysosomal Pathway. Front. Cell Dev. Biol. 2020, 8, 602574, 10.3389/fcell.2020.602574.

29. Liao, Z.; Chen, Y.; Zhang, Y.; Li, Y.; Li, J.; Li, X.; Zhao, L.; Zhang, L.; Chen, G. Liraglutide Improves Nonalcoholic Fatty Liver Disease in Diabetic Mice by Activating Autophagy Through AMPK/mTOR Signaling Pathway. Diabetes Metab. Syndr. Obes. 2024, 17, 643–656, 10.2147/DMSO.S447182.

30. Yang, M.; Zhang, X.; Li, X.; Wang, X.; Deng, Y.; Li, Y.; Wang, Y.; Li, J.; Zhang, Z.; Wang, Z. Liraglutide Attenuates Non-Alcoholic Fatty Liver Disease in Mice by Regulating the Local Renin Angiotensin System. Front. Pharmacol. 2020, 11, 432, 10.3389/fphar.2020.00432.

31. Armstrong, M.J.; Gaunt, P.; Aithal, G.P.; Barton, D.; Hull, D.; Parker, R.; Hazlehurst, J.M.; Guo, K.; Abouda, G.; Aldersley, M.A.; et al. Liraglutide safety and efficacy in patients with non alcoholic steatohepatitis (LEAN): a multicentre, double-blind, randomised, placebo-controlled phase 2 study. Lancet 2016, 387, 679–690, 10.1016/S0140-6736(15)00803-X.

32. Tan, Y.; Zhang, Y.; Ling, X.; Lu, Y.; Chen, Y.; Qin, X.; Li, X. Association between use of liraglutide and liver fibrosis in patients with type 2 diabetes. Front. Endocrinol. 2022, 13, 935180, 10.3389/fendo.2022.935180.

33. Ji, J.; Yang, L.; He, J.; Wang, Z. Liraglutide inhibits receptor for advanced glycation end products (RAGE)/reduced form of nicotinamide-adenine dinucleotide phosphate (NAPDH) signaling to ameliorate non-alcoholic fatty liver disease (NAFLD) in vivo and vitro. Bioengineered 2022, 13, 5723–5737, 10.1080/21655979.2022.2036902.

34. Fasano, T.; Pisciotta, L.; Bocchi, L.; Guardamagna, O.; Assandro, P.; Rabacchi, C.; Zanoni, P.; Filocamo, M.; Bertolini, S.; Calandra, S. Lysosomal lipase deficiency: molecular characterization of eleven patients with wolman or cholesteryl ester storage disease. Mol. Genet. Metab. 2012, 105, 450–456, 10.1016/j.ymgme.2011.12.008.

35. Tovoli, F.; Napoli, L.; Negrini, G.; D’Addato, S.; Tozzi, G.; D’Amico, J.; Piscaglia, F.; Bolondi, L. A Relative Deficiency of Lysosomal Acid Lypase Activity Characterizes Non-Alcoholic Fatty Liver Disease. Int. J. Mol. Sci. 2017, 18, 1134, 10.3390/ijms18061134.

36. Shteyer, E.; Villenchik, R.; Mahamid, M.; Nator, N.; Safadi, R. Low Serum Lysosomal Acid Lipase Activity Correlates with Advanced Liver Disease. Int. J. Mol. Sci. 2016, 17, 312, 10.3390/ijms17030312.

37. Gomaraschi, M.; Bonacina, F.; Norata, G.D. Lipid accumulation impairs lysosomal acid lipase activity in hepatocytes: evidence in nafld patients and cell cultures. Biochim. Biophys. Acta Mol. Cell Biol. Lipids 2019, 1864, 158523, 10.1016/j.bbalip.2019.158523.

38. Leopold, C.; Duta-Mare, M.; Sachdev, V.; Goeritzer, M.; Maresch, L.K.; Kolb, D.; Reicher, H.; Wagner, B.; Stojakovic, T.; Ruelicke, T.; et al. Hepatocyte-specific lysosomal acid lipase deficiency protects mice from diet-induced obesity but promotes hepatic inflammation. Biochim. Biophys. Acta Mol. Cell Biol. Lipids 2019, 1864, 500–511, 10.1016/j.bbalip.2019.01.007.

39. Lopresti, M.W.; Ahn, J.H.; Wencewicz, T.A.; Farese, R.V.; Walther, T.C. Hepatic lysosomal acid lipase overexpression worsens hepatic inflammation in mice fed a Western diet. J. Lipid Res. 2021, 62, 100133, 10.1016/j.jlr.2021.100133.

40. Mizushima, N.; Yamamoto, A.; Matsui, M.; Yoshimori, T.; Ohsumi, Y. In Vivo Analysis of Autophagy in Response to Nutrient Starvation Using Transgenic Mice Expressing a Fluorescent Autophagosome marker. Mol. Biol. Cell 2004, 15, 1101–1111, 10.1091/mbc.e03-09-0704.

41. Martinez-Lopez, N.; Tarabra, E.; Toledo, M.; Garcia-Macia, M.; Sahu, S.; Coletto, L.; Batista Gonzalez, A.; Barzilai, N.; Pessin, J.E.; Schwartz, G.J.; et al. System-wide Benefits of Intermeal Fasting by Autophagy. Cell Metab. 2017, 26, 856–871, 10.1016/j.cmet.2017.09.020.

42. Moreira, G.V.; Azevedo, F.F.; Ribeiro, L.M.; Santos, A.; Guadagnini, D.; Gama, P.; Carvalho, C.R.O. Liraglutide modulates gut microbiota and reduces NAFLD in obese mice. J. Nutr. Biochem. 2018, 62, 143–154, 10.1016/j.jnutbio.2018.07.009.

43. Sturis, J.; Gotfredsen, C.F.; Rømer, J.; Rolin, B.; Ribel, U.; Brand, C.L.; Wilken, M.; Wassermann, K.; Deacon, C.F.; Carr, R.D.; et al. GLP-1 derivative liraglutide in rats with β-cell deficiencies: influence of metabolic state on β-cell mass dynamics. Br. J. Pharmacol. 2003, 140, 123–132, 10.1038/sj.bjp.0705397.

44. Wang, Y.; Ji, B.; Wu, W.; Wang, R.; Yang, Z.; Zhang, D.; Tian, W. Hepatoprotective effects of kombucha tea: identification of functional strains and quantification of functional components. J. Sci. Food Agric. 2013, 94, 265–272, 10.1002/jsfa.6245.

45. Silva, F.F.; Monteiro, L.G.; Trombetta, I.C.; Oliveira, E.P.; de Fante, T.; de Paula, F.J.A.; de Sá, J.C.F.; de Oliveira, M.A.M.; de Carvalho, P.O.; de Almeida-Pittito, B. Dexamethasone-Induced Adipose Tissue Redistribution and Metabolic Changes: is gene expression the main factor? an animal model of chronic hypercortisolism. Biomedicines 2022, 10, 2328, 10.3390/biomedicines10092328.

46. Agouni, A.; Tual-Chalot, S.; Chalopin, M.; Duluc, L.; Mody, N.; Schwarz, P.E.H.; Withers, D.J.; Martinez, M.C.; Andriantsitohaina, R. In vivo differential effects of fasting, re-feeding, insulin and insulin stimulation time course on insulin signaling pathway components in peripheral tissues. Biochem. Biophys. Res. Commun. 2010, 401, 104–111, 10.1016/j.bbrc.2010.09.018.

47. Andrikopoulos, S.; Blair, A.R.; Deluca, N.; Fam, B.C.; Proietto, J. Evaluating the glucose tolerance test in mice. Am. J. Physiol. Endocrinol. Metab. 2008, 295, E1323–E1332, 10.1152/ajpendo.90617.2008.

48. Ahmed, L.A.; Obaid, A.A.; Zaki, H.F.; Agha, A.M. Gut microbiota modulation as a promising therapy with metformin in rats with non-alcoholic steatohepatitis: role of lps/tlr4 and autophagy pathways. Eur. J. Pharmacol. 2020, 887, 173461, 10.1016/j.ejphar.2020.173461.

49. Martinet, W.; Agostinis, P.; Vanhoecke, B.; Dewaele, M.; De Meyer, G.R.Y. Immunohistochemical analysis of macroautophagy. Autophagy 2013, 9, 386–402, 10.4161/auto.22968.

50. Friedewald, W.T.; Levy, R.I.; Fredrickson, D.S. Estimation of the concentration of low-density lipoprotein cholesterol in plasma, without use of the preparative ultracentrifuge. Clin. Chem. 1972, 18, 499–502. Available online: https://pubmed.ncbi.nlm.nih.gov/4337382/.

51. Araujo, L.C.C.; Feitosa, K.B.; Murata, G.M.; Furigo, I.C.; Teodoro, B.G.; de Souza, C.O.; dos Santos, G.A.; de Sá, R.C.C.; Solon, C.; de Bernardes, S.S.; et al. Uncaria tomentosa improves insulin sensitivity and inflammation in experimental NAFLD. Sci. Rep. 2018, 8, 11013, 10.1038/s41598-018-29044-y.

52. Mehlem, A.; Hagberg, C.E.; Muhl, L.; Eriksson, U.; Falkevall, A. Imaging of neutral lipids by oil red O for analyzing the metabolic status in health and disease. Nat. Protoc. 2013, 8, 1149–1154, 10.1038/nprot.2013.055.

53. Ramírez-Zacarías, J.L.; Castro-Muñozledo, F.; Kuri-Harcuch, W. Quantitation of adipose conversion and triglycerides by staining intracytoplasmic lipids with oil red O. Histochemistry 1992, 97, 493–497, 10.1007/BF00316069.

54. Folch, J.; Lees, M.; Sloane Stanley, G.H. A simple method for the isolation and purification of total lipides from animal tissues. J. Biol. Chem. 1957, 226, 497–509. Available online: https://pubmed.ncbi.nlm.nih.gov/13428781/.

