## Supplemental Figures S1-S6 and supplemental Table S1 for "Liraglutide exerts protective effects on MASLD independently of lipophagy, and the combination with kombucha presents no additional advantage"

**Supplementary Figure S1.** Body weight and composition of control animals and those fed a high-fat diet.

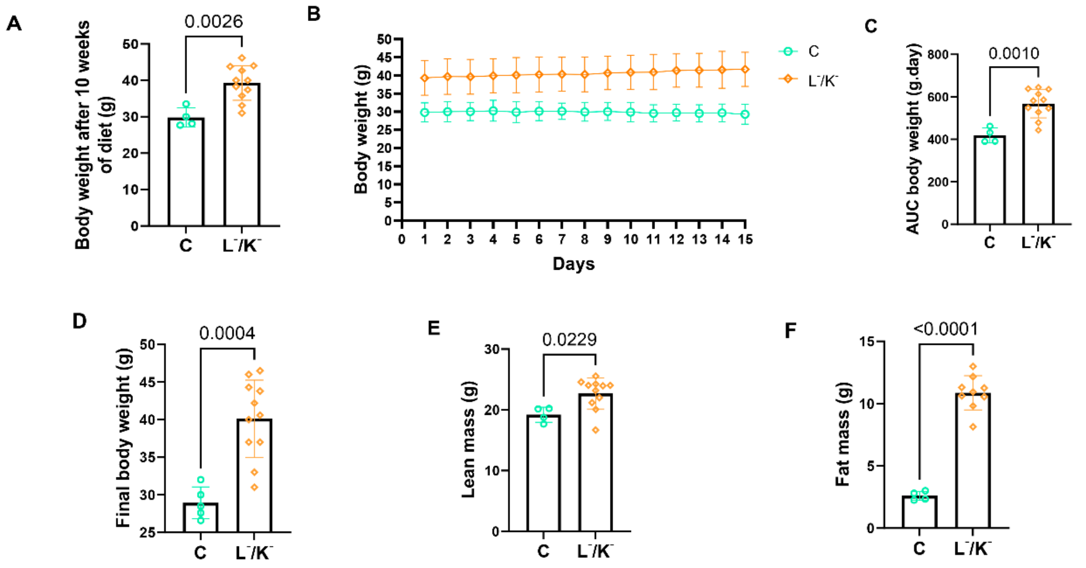


**(A)** Body weight after 10 weeks of diet (n = 4 in C and n = 11 in L-/K-). **(B)** Time course of body weight from the 1st to the 15th day of treatment (n = 4 in C and n = 11 in L-/K-). **(C)** Area under the curve (AUC) of body weight of animals throughout the treatment period (n = 4 in C and n = 11 in L-/K-). **(D)** Final body weight, obtained after the last day of the treatment period (n = 5 in C and n = 11 in L-/K-). **(E)** Lean body mass content, obtained after the last day of the treatment period (n = 4 in C and n = 11 in L-/K-). **(F)** Fat body mass content, obtained after the last day of the treatment period (n = 4 in C and n = 9 in L-/K-). Each scatter plot indicates a sampling unit. The results were obtained through Student's t-test, are presented as MD ± SD and are indicated by P value. The significance level adopted was $P \leq 0.05$.

**Supplementary Table S1.** Food and caloric intake of control animals and those fed a high-fat diet.

| Group | AUC Food intake (g.day) Mean ± SD | AUC Caloric intake (kJ.day) Mean ± SD |
|---|---|---|
| C | 50.608 ± 1.671 | 677.560 ± 22.335 |
| $L^-/K^-$ | 38.195 ± 4.524 *** | 854.933 ± 101.328 ** |

AUC of food intake during the treatment period (n = 5 in C and n = 6 in the L-/K- group). AUC of caloric intake throughout the treatment period (n = 5 in C and n = 6 in the L-/K- group). The results were obtained through Student's t-test, are presented as MD ± SD, and the significance level adopted was P ≤ 0.05. *** indicates P ≤ 0.001 in relation to C, and ** indicates P ≤ 0.01 in relation to C.

**Supplementary Figure S2.** Fasting blood glucose and glucose tolerance of control animals and those fed a high-fat diet.

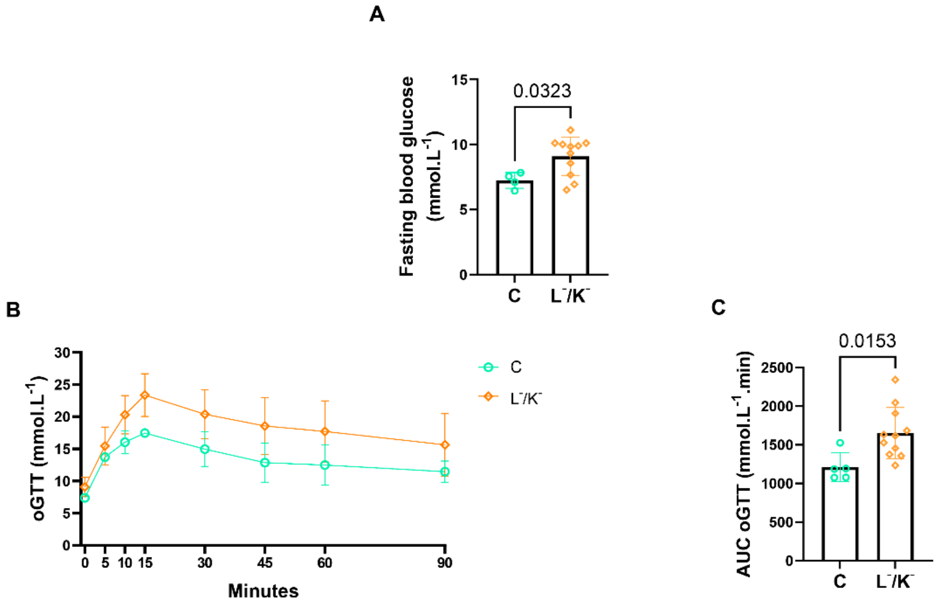


**(A)** Fasting blood glucose levels obtained at the end of the treatment period (n = 4 in C and n = 11 in L-/K-). **(B)** Time course of the oral glucose tolerance test (oGTT) performed at the end of the treatment period (n = 5 in C and n = 11 in L-/K-). **(C)** AUC of the oGTT (n = 5 in C and n = 11 in L-/K-). Each scatter plot indicates a sampling unit. The results were obtained through Student's t-test, are presented as MD ± SD and are indicated by P value. The significance level adopted was $P \leq 0.05$.

**Supplementary Figure S3.** Plasma lipids of control animals and those fed a high-fat diet.

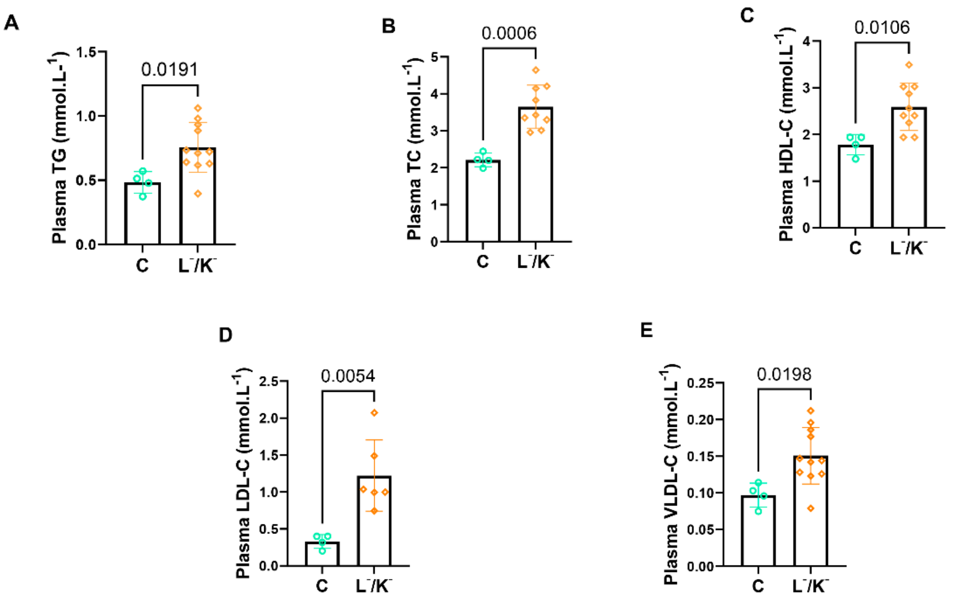


**(A)** Plasma triglyceride (TG) levels (n = 4 in C and n = 11 in L-/K-) **(B)** Plasma total cholesterol (TC) levels (n = 4 in C and n = 9 in L-/K-) **(C)** Plasma high-density lipoprotein cholesterol (HDL-C) levels (n = 4 in C and n = 10 in L-/K-). **(D)** Plasma low-density lipoprotein cholesterol (LDL-C) levels (n = 4 in C and n = 6 in L-/K-). **(E)** Plasma very low-density lipoprotein cholesterol (VLDL-C) levels (n = 4 in C and n = 11 in L-/K-). Each scatter plot indicates a sampling unit. The results were obtained through Student's t-test, are presented as MD ± SD and are indicated by P value. The significance level adopted was $P \leq 0.05$.

**Supplementary Figure S4.** Hepatic histology of control animals and those fed a high-fat diet.

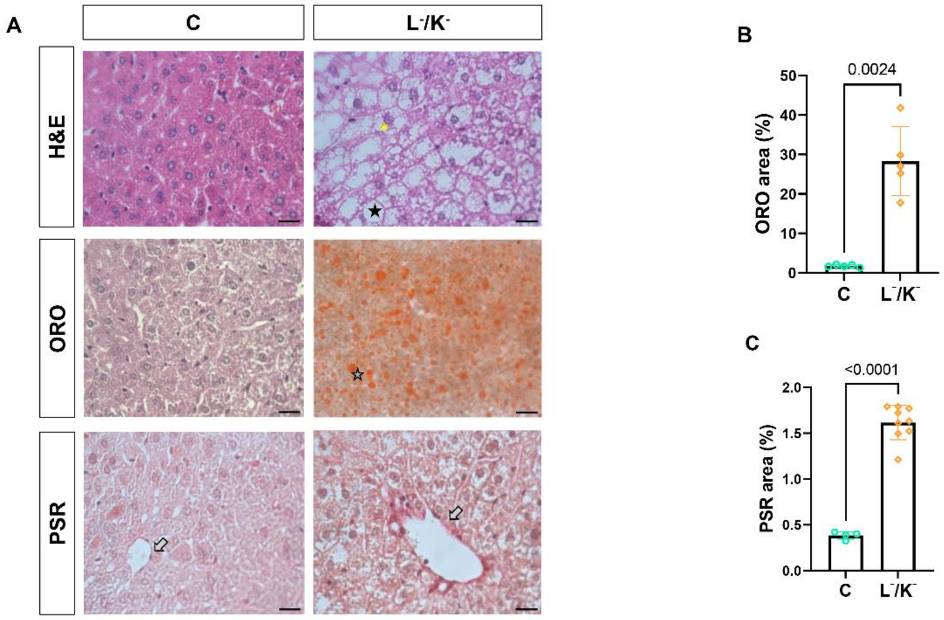


**(A)** Representative images of Hematoxylin and Eosin (H&E), Oil Red O (ORO), and Picrosirius Red (PSR) staining. **(B)** liver area stained by ORO staining (n = 5 in C and L-/K-). **(C)** liver area stained by PSR staining (n = 4 in C and n = 9 in L-/K-). Each scatter plot indicates a sampling unit. The results were obtained through Student's t-test, are presented as MD ± SD and are indicated by P value. The significance level adopted was $P \leq 0.05$. Magnification of 60× and scale bar of 17 μm in H&E, PSR, and ORO. Black star indicates the presence of vesicles, yellow asterisk indicates the presence of hepatocellular ballooning, gray star indicates the accumulation of neutral lipids, and gray arrows indicate the presence of collagen fibers in the perivascular area.

**Supplementary Figure S5.** Hepatic parameters of control animals and those fed a high-fat diet.

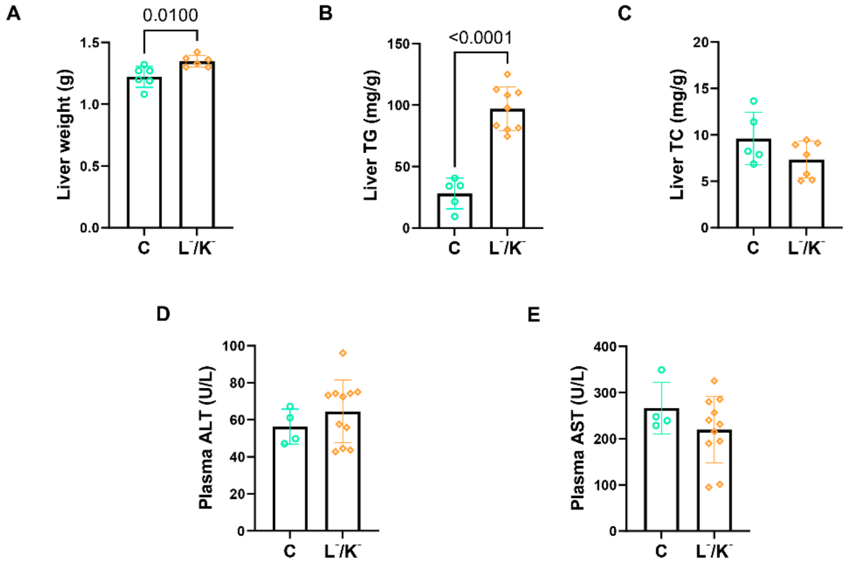


**(A)** Liver weight (n = 6 in C and L-/K-) **(B)** Hepatic TG content (n = 5 in C and n = 9 in L-/K-). **(C)** Hepatic TC content (n = 5 in C and n = 7 in L-/K-). **(D)** Plasma alanine aminotransferase (ALT) levels (n = 4 in C and n = 11 in L-/K-). **(E)** Plasma aspartate aminotransferase (AST) levels (n = 4 in C and n = 11 in L-/K-). Each scatter plot indicates a sampling unit. The results were obtained through Student's t-test, are presented as MD ± SD and are indicated by P value. The significance level adopted was $P \le 0.05$.

**Supplementary Figure S6.** Hepatic autophagy-related proteins and lysosomal acid lipase (LAL) expression and activity in control animals and those fed a high-fat diet.

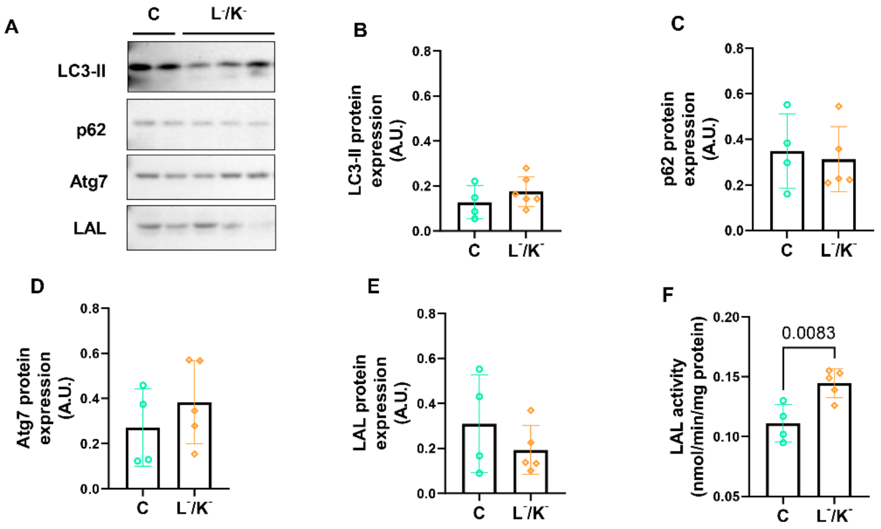


**(A)** Representative image of hepatic expression of Microtubule-Associated Protein 1A/1B-Light Chain 3 (LC3-II), Sequestosome-1 (p62), Autophagy Related 7 (Atg7), and LAL. **(B)** Hepatic expression of LC3-II, obtained by Western blotting (n = 4 in C and n = 6 in L-/K-). **(C)** Hepatic expression of p62, obtained by Western blotting (n = 4 in C and n = 5 in L-/K-). **(D)** Hepatic expression of Atg7, obtained by Western blotting (n = 4 in C and n = 5 in L-/K-). **(E)** Hepatic expression of LAL, obtained by Western blotting (n = 4 in C and n = 5 in L-/K-). **(F)** LAL activity in the liver, obtained by fluorescence assay (n = 4 in C and n = 5 in L-/K-). Two independent membranes were used for statistical analysis (total n = 4–6 per group). Image representing all groups (C, L-/K-, L+/K-, L-/K+, and L+/K+), cropped for representative purposes in this figure. The same samples from the L-/K- group were used in both this figure and Figure 6, A. Each scatter plot indicates a sampling unit. The results were obtained through Student's t-test, are presented as MD ± SD and are indicated by P value. The significance level adopted was $P \leq 0.05$.
